# Autophagosome-tethering compounds selectively lower mutant ataxin-3 in preclinical models of spinocerebellar ataxia type 3

**DOI:** 10.64898/2026.10.02.756117

**Authors:** Hongyu Yuan, Zhao Chen, Xiaokai Shen, Yiqing Gong, Yan Tan, Jiawei He, Na Wan, Chunrong Wang, Linlin Wan, Linliu Peng, Lang He, Huirong Peng, Rong Qiu, Beisha Tang, Boxun Lu, Hong Jiang

## Abstract

Spinocerebellar ataxia type 3 (SCA3) is most common subtype of autosomal dominant spinocerebellar ataxia worldwide and the most prevalent CAG expansion disorder in the east Asian population, but disease-modifying therapies remain unavailable. SCA3 is caused by the accumulation of polyglutamine- (polyQ-) expanded mutant ataxin-3 (mAtxn3), while the wild-type ataxin-3 (wtAtxn3) plays an important cellular role. Thus, allele-selective clearance of mAtxn3 without affecting wtAtxn3 is highly desired for SCA3 treatment. To achieve this goal, we tested previously reported compounds that have been shown to selectively degrade the Huntington’s disease causing mutant HTT protein by tethering the expanded polyQ to the autophagosome protein LC3. Here, we demonstrated that two of these compounds, AN2 and GW5074, selectively reduced both soluble and aggregated mAtxn3 with expanded polyQ (84Q), but not wtAtxn3, in both cellular and mouse models of SCA3. Mechanistically, both compounds lowered mAtxn3 by enhancing its autophagic degradation without obvious activation of global autophagy. At the molecular level, both compounds increased mAtxn3-LC3 interaction and colocalization to enhance the allele-selectively autophagic degradation of mAtxn3. Phenotypically, both compounds rescued cell viability and ATP content of SCA3 cellular models. In SCA3 patient pluripotent stem cells (iPSCs) derived cortical neurons, both compounds allele-selectively reduced mAtxn3 and partially rescued neurite shortening. Importantly, in a well-established SCA3 transgenic mouse model (MJD84.2), intracerebroventricular, intraperitoneal, and oral administration allele-selectively reduced mAtxn3 in multiple brain regions and rescued motor function deficits. Oral dosing achieved nanomolar cerebellar exposure and showed no overt liver, kidney, or glial toxicity over 28 days. Analyses of 1303 Chinese patients with SCA3 and ATXN3 constructs of different polyglutamine lengths further supported the compounds’ polyQ length-dependent activity with relative sparing of normal-length ataxin-3. Together, these findings provide preclinical proof of concept that utilizing orally available small molecule compounds to allele-selectively enhance mAtxn3 autophagic degradation for SCA3 treatment.

**One-sentence summary:** Small-molecule autophagosome-tethering compounds selectively lower mutant ataxin-3 through an LC3- and autophagy-dependent mechanism and improve disease-related phenotypes in preclinical models of SCA3.

## Introduction

Spinocerebellar ataxia type 3 (SCA3), also known as Machado-Joseph disease, is the most common subtype of autosomal dominant spinocerebellar ataxia worldwide and is highly represented in Chinese ataxia cohorts [1–3]. SCA3 is caused by CAG-repeat expansion in the coding region of gene *ATXN3*, which generates ataxin-3 with an expanded polyglutamine (polyQ) tract [4–6]. The mutant protein misfolds, accumulates in neuronal nuclei, and forms aggregation-prone species in vulnerable regions such as the deep cerebellar nuclei and pons [5–7]. Despite slow clinical progression, SCA3 remains relentlessly disabling, and no disease-modifying therapy is available.

Therapeutic strategies for SCA3 have focused on reducing mutant *ATXN3* expression or enhancing clearance of mutant ataxin-3, including antisense oligonucleotides, RNA interference, autophagy induction, and proteostasis-modulating small molecules [8–11]. Although nucleic-acid approaches are mechanistically close to the genetic cause of SCA3, their clinical development must address central nervous system delivery, relatively high immunogenicity, allele selectivity, durability, and long-term safety [12–14]. Importantly, these gene therapy approaches are not orally available, preventing patients from taking medicines at home. Thus, direct clearance of the already synthesized pathogenic protein therefore represents a complementary therapeutic strategy.

Autophagy-based targeted protein degradation is well suited to polyQ diseases because autophagy can eliminate aggregation-prone and insoluble protein species that may be poor proteasomal substrates [15]. Li et al. previously identified HTT–LC3 linker compounds, later referred to as autophagosome-tethering compounds (ATTECs), that interact with both LC3 and expanded polyQ-containing mutant huntingtin, promote autophagic degradation of mutant huntingtin, and ameliorate disease phenotypes in Huntington disease models [9, 16]. This work suggested that ATTECs may recognize conformational features associated with expanded polyQ tracts rather than mutant huntingtin alone, raising the possibility of extending this strategy to other polyQ disorders. Consistent with this idea, preliminary data suggests that they may reduce mAtxn3 in a single SCA3 patient-derived fibroblast line[9], and related indole and coumarin derivatives enhanced the interaction of mutant ATXN3-Q75 with LC3 and mitigated aggregation-associated phenotypes in an ATXN3-Q75 cell model [17]. However, ATTEC-based targeting of ataxin-3 has remained largely limited to preliminary cellular evidence, and its mechanism, in vivo efficacy, tolerability, oral availability, activity in patient-derived neurons, and selectivity over normal-length ataxin-3, and importantly, therapeutic potentials, have not been systematically tested. Such information is of vital importance for the development of potential SCA3 therapeutic based on this strategy. Mechanistically, validation of these compounds in vivo is also important to evaluate the pathogenic contributions of the mAtxn3 protein alone without influencing the gene sequence or mRNA expression. This is especially important in light of recent studies showing that changes at the DNA or mRNA level, such as somatic expansion or phase-separation, may also contribute to the disease [18, 19].

Thus, we evaluated AN2 and GW5074, two previously reported ATTECs that reduced mHTT levels by engaging expanded polyQ [9] for their effects on the polyQ-expanded mAtxn3 and SCA3-relevant disease phenotypes. We assessed their selectivity, effects on aggregation-prone ataxin-3 species, LC3- and autophagy-dependent mechanism, and therapeutic activity across intracerebroventricular, intraperitoneal, and oral delivery routes. Importantly, we also assessed oral brain exposure and tolerability by orally administrations, suggesting therapeutic potentials. Finally, we combined a 1303-patient Chinese SCA3 cohort with ATXN3 constructs spanning different polyQ lengths to assess the relationship between repeat length and compound response.

## Results

### AN2 and GW5074 selectively lowered mAtxn-3 in cellular SCA3 models

We first tested AN2 and GW5074 in HEK293T cells transiently expressing FLAG-tagged ATXN3-20Q or ATXN3-84Q. After 48 hours of treatment with DMSO or compounds at 30, 50, 100, 200, or 300nM, neither compound detectably altered the wild-type 20Q ataxin-3, whereas both reduced the disease-causing 84Q ataxin-3, with the strongest effect at 100nM and slightly weaker lowering at 300nM. This bell-shaped response was consistent with previous reports [9] and was accompanied by partial restoration of the reduced cell viability and ATP content associated with ATXN3-84Q expression (Supplementary Fig. 1). Similar allele-selective lowering and cell viability improvement were observed in SH-SY5Y stable expression models, supporting activity in a neuronal-lineage cellular context (Fig. 1). Control immunoblots using anti-FLAG and anti-polyQ antibodies, together with whole-membrane blotting, excluded antibody-epitope bias or enhanced site-specific cleavage as the main explanation for the reduced full-length 84Q ataxin-3 signal (Supplementary Fig. 2). Together, these results indicate that AN2 and GW5074 selectively lowered mAtxn3 with expanded polyQ ataxin-3 and rescued cellular impairment in two cellular SCA3 models.

**Fig. 1.**
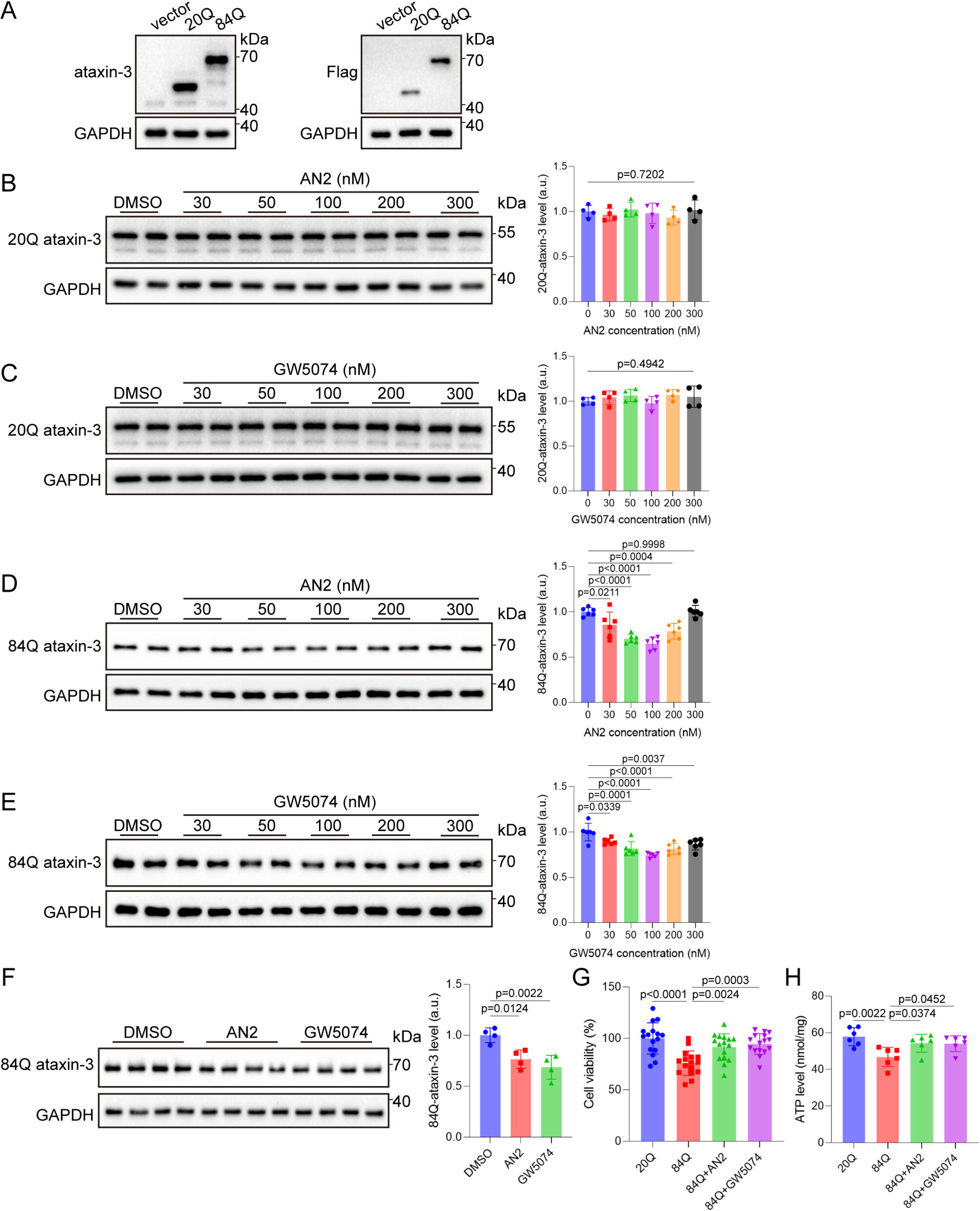
AN2 and GW5074 selectively reduce mutant ataxin-3 and improve cellular phenotypes in SH-SY5Y stable expression models. **(A)** Representative immunoblots of SH-SY5Y cells stably expressing empty vector, ATXN3-20Q-FLAG, or ATXN3-84Q-FLAG, showing specific ataxin-3 and FLAG signals. **(B to E)** Representative immunoblots and quantification of 20Q ataxin-3 **(B and C)** or 84Q ataxin-3 **(D and E)** in SH-SY5Y stable expression models treated with DMSO or the indicated concentrations of AN2 or GW5074 for 48 hours. Protein abundance was normalized to GAPDH and then to the mean value of the DMSO group. Data are shown as mean ± SD; n = 4 for **(B)** and **(C)**, and n = 6 for **(D)** and **(E)**. **(F)** Representative immunoblot and quantification of 84Q ataxin-3 in SH-SY5Y cells stably expressing ATXN3-84Q-FLAG after treatment with DMSO, AN2 (100nM), or GW5074 (100nM) for 48 hours. Protein abundance was normalized to GAPDH and then to the mean value of the DMSO group. Data are shown as mean ± SD; n = 4. For **(B) to (F)**, statistical analysis was performed by one-way ANOVA followed by Dunnett’s multiple-comparisons test versus the DMSO group. **(G)** Cell viability of SH-SY5Y cells stably expressing 20Q or 84Q ataxin-3 after treatment with DMSO, AN2 (100nM), or GW5074 (100nM). Values were normalized to the mean value of the 20Q ataxin-3 group and expressed as percentages. Data are shown as mean ± SD; n = 16. **(H)** ATP content in SH-SY5Y cells stably expressing 20Q or 84Q ataxin-3 after treatment with DMSO, AN2 (100nM), or GW5074 (100nM). Data are shown as mean ± SD; n = 6. For **(G)** and **(H)**, statistical analysis was performed by one-way ANOVA followed by Dunnett’s multiple-comparisons test versus the 84Q ataxin-3 DMSO group. *P* values are shown in the plots.

### AN2 and GW5074 preferentially reduced aggregation-associated mutant ataxin-3 species

We next examined whether AN2 and GW5074 preferentially affected aggregation-associated mutant ataxin-3 species. Sequential extraction with 1% Triton X-100 showed that both compounds reduced 84Q ataxin-3 in soluble and insoluble fractions, with a stronger effect in the insoluble fraction: AN2 and GW5074 reduced soluble 84Q ataxin-3 to 78.95% and 76.34% of DMSO, respectively, but reduced insoluble 84Q ataxin-3 to 55.28% and 49.51% (Fig. 2, A and B). Using EGFP-tagged ATXN3 constructs, we found that EGFP-ATXN3-84Q, but not EGFP-ATXN3-20Q, generated high-molecular-weight SDS-resistant signals mainly in the Triton X-100-insoluble fraction by days 5 to 7; at day 7, AN2 and GW5074 reduced these SDS-resistant mutant ataxin-3 signals (Supplementary Fig. 3, A and B; Fig. 2, C to E). Consistent with the immunoblot data, confocal imaging showed that both compounds reduced the proportion of aggregate-positive EGFP-Ataxin3Q84-expressing HeLa cells (Fig. 2, F and G). Together, these biochemical and imaging results indicate that AN2 and GW5074 preferentially reduced aggregation-associated mutant ataxin-3 species.

**Fig. 2.**
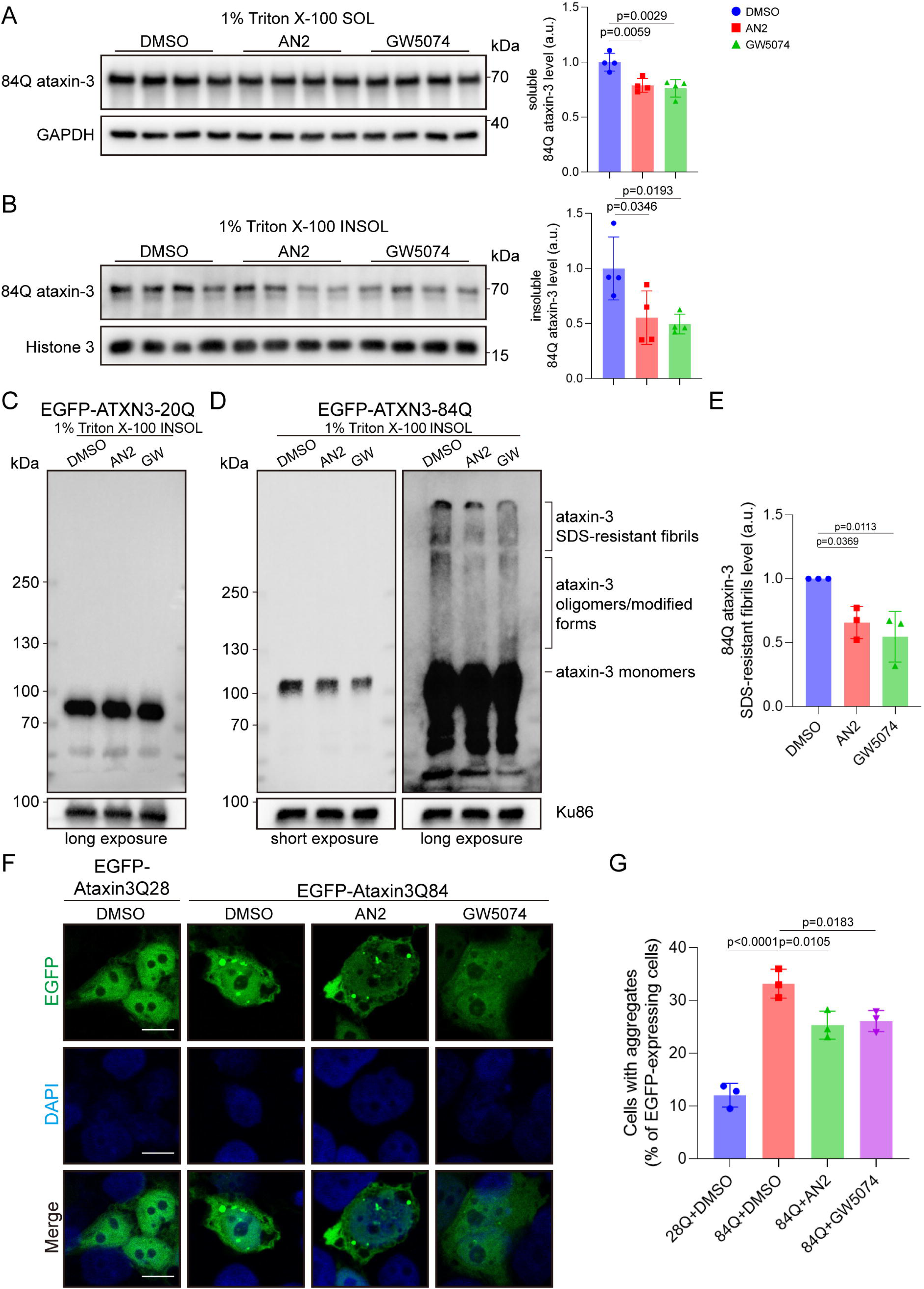
AN2 and GW5074 preferentially reduce insoluble and aggregation-associated mutant ataxin-3 species. **(A and B)** Representative immunoblots and quantification of 84Q ataxin-3 in the 1% Triton X-100-soluble fraction **(A)** and Triton X-100-insoluble fraction **(B)** from HEK293T cells transiently expressing ATXN3-84Q-FLAG after treatment with DMSO, AN2 (100nM), or GW5074 (100nM) for 48 hours. Protein abundance was normalized to the corresponding loading control and then to the mean value of the DMSO group. Data are shown as mean ± SD; n = 4. Statistical analysis was performed by one-way ANOVA followed by Dunnett’s multiple-comparisons test versus the DMSO group. *P* values are shown in the plots. SOL, 1% Triton X-100-soluble fraction; INSOL, 1% Triton X-100-insoluble fraction. **(C)** Representative long-exposure immunoblot of the 1% Triton X-100-insoluble fraction from HEK293T cells transiently expressing EGFP-ATXN3-20Q after treatment with DMSO, AN2 (100nM), or GW5074 (100nM) for 7 days. **(D and E)** Representative short-exposure and long-exposure immunoblots **(D)** and quantification of SDS-resistant ataxin-3 fibril signals **(E)** in the 1% Triton X-100-insoluble fraction from HEK293T cells transiently expressing EGFP-ATXN3-84Q after treatment with DMSO, AN2 (100nM), or GW5074 (100nM) for 7 days. Ku86 was used as the reference protein for normalization of the insoluble fraction, and the normalized values were further expressed relative to the mean value of the DMSO group. Data are shown as mean ± SD; n = 3. Statistical analysis was performed by one-way ANOVA followed by Dunnett’s multiple-comparisons test versus the DMSO group. *P* values are shown in the plot. INSOL, 1% Triton X-100-insoluble fraction; GW, GW5074. **(F and G)** Representative confocal images **(F)** and quantification of aggregate-positive cells **(G)** in HeLa cells transiently expressing EGFP-Ataxin3Q28 or EGFP-Ataxin3Q84 after treatment with DMSO, AN2 (100nM), or GW5074 (100nM). EGFP-positive ataxin-3 aggregates were quantified as the percentage of EGFP-expressing cells containing visible aggregates. Images were obtained from three independent experiments, with 20 fields analyzed per experiment. Data are shown as mean ± SD; n = 3 independent experiments. Statistical analysis was performed by one-way ANOVA followed by Dunnett’s multiple-comparisons test versus the EGFP-Ataxin3Q84 DMSO group. *P* values are shown in the plot. Scale bars, 10μm.

### AN2 and GW5074 enhanced mAtxn3-LC3 proximity and enhanced autophagy-dependent mAtxn-3 degradation

To test whether AN2 and GW5074 promote mutant ataxin-3-LC3 association in cells, we used validated HA-TurboID-tagged ATXN3 proximity-labeling models in HEK293T cells (Supplementary Fig. 3, C and D). AN2 and GW5074 increased the LC3/84Q ataxin-3 ratio in streptavidin-enriched fractions, whereas no comparable increase was detected for 20Q ataxin-3; comparable streptavidin-HRP patterns among biotin-treated groups excluded the possibility that the increase was not due to nonspecific differences in labeling efficiency (Fig. 3, A to F). At the cellular level, confocal imaging of HeLa cells coexpressing EGFP-Ataxin3Q84 and RFP-LC3 showed increased LC3-colocalized ataxin-3 puncta after compound treatment, whereas EGFP-Ataxin3Q28 showed little colocalized punctate signal (Fig. 3, G and H).

**Fig. 3.**
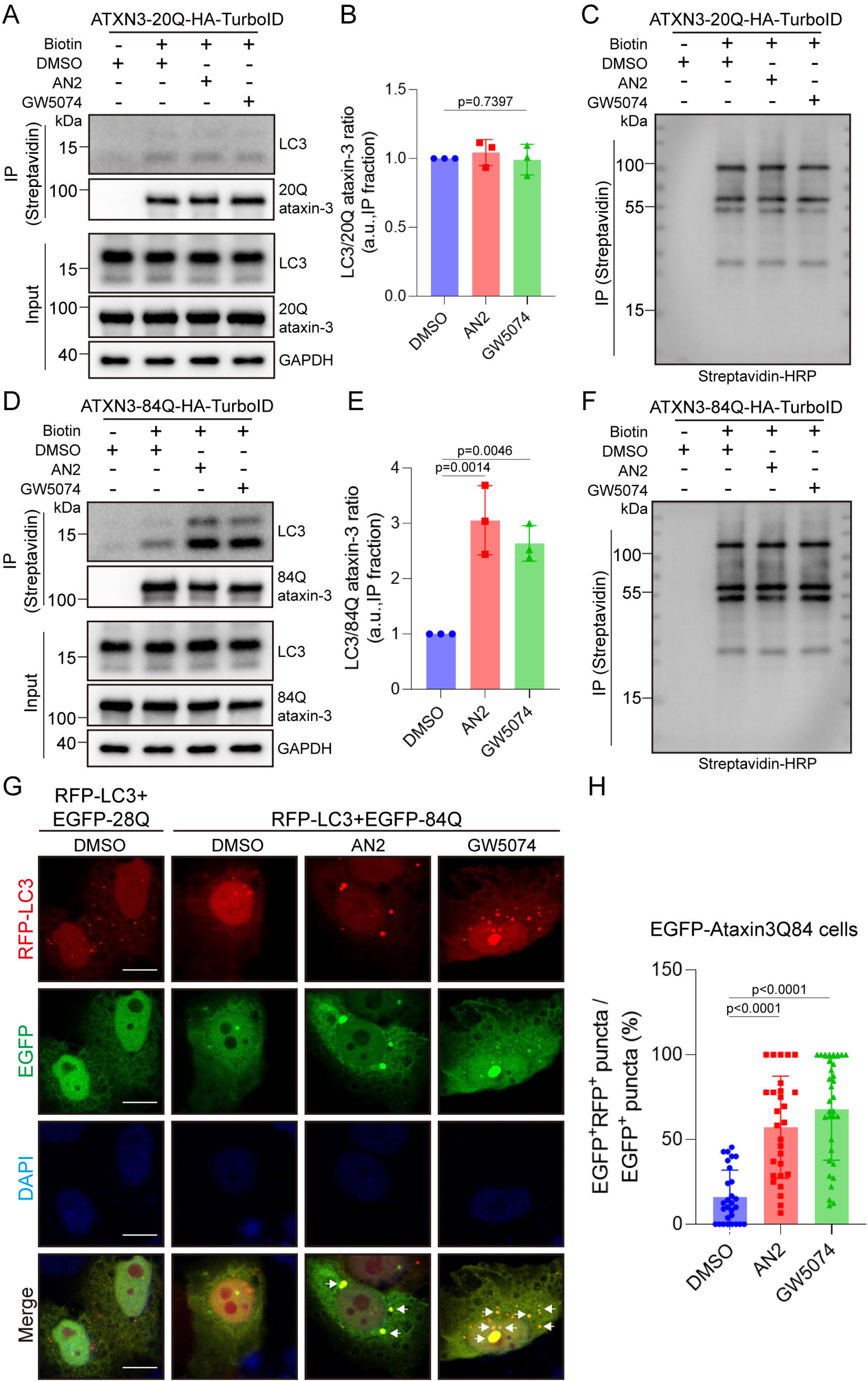
AN2 and GW5074 increase LC3 proximity and colocalization with mutant ataxin-3. **(A)** Representative immunoblots of streptavidin-enriched fractions and input lysates from HEK293T cells expressing ATXN3-20Q-HA-TurboID. Cells were treated with or without 100μM biotin and with DMSO, AN2 (100nM), or GW5074 (100nM) for 4 hours. LC3 and 20Q ataxin-3 were detected in streptavidin-enriched fractions and input lysates; GAPDH was used as a loading control for input samples. **(B)** Quantification of the LC3/20Q ataxin-3 ratio in streptavidin-enriched fractions from **(A)**. Values were normalized to the mean value of the biotin-positive DMSO group. **(C)** Streptavidin-HRP blot of streptavidin-enriched fractions from the same experimental conditions as in **(A)**. **(D)** Representative immunoblots of streptavidin-enriched fractions and input lysates from HEK293T cells expressing ATXN3-84Q-HA-TurboID and treated as in **(A)**. LC3 and 84Q ataxin-3 were detected in streptavidin-enriched fractions and input lysates; GAPDH was used as a loading control for input samples. **(E)** Quantification of the LC3/84Q ataxin-3 ratio in streptavidin-enriched fractions from **(D)**. Values were normalized to the mean value of the biotin-positive DMSO group. **(F)** Streptavidin-HRP blot of streptavidin-enriched fractions from the same experimental conditions as in **(D)**. For **(B)** and **(E)**, data are shown as mean ± SD; n = 3. Statistical analysis was performed by one-way ANOVA followed by Dunnett’s multiple-comparisons test versus the DMSO group. **(G)** Representative confocal images of HeLa cells cotransfected with RFP-LC3 and EGFP-Ataxin3Q28 or EGFP-Ataxin3Q84 and treated with DMSO, AN2 (100nM), or GW5074 (100nM) for 4 hours. Green, EGFP-ataxin-3; red, RFP-LC3; blue, DAPI. White arrows indicate representative EGFP- and RFP-double-positive puncta. Scale bars, 10 μm. **(H)** Quantification of colocalized puncta in EGFP-Ataxin3Q84-expressing cells, calculated as the percentage of EGFP- and RFP-double-positive puncta among total EGFP-positive puncta. Data are shown as mean ± SD; n = 28 to 33. Statistical analysis was performed by Welch’s one-way ANOVA followed by Dunnett’s T3 multiple-comparisons test versus the DMSO group. *P* values are shown in the plots.

In SH-SY5Y cells stably expressing ATXN3-84Q-FLAG, pretreatment with bafilomycin A1 (BafA1, 100nM) or chloroquine (CQ, 40μM) abolished the lowering effect of AN2 and GW5074 on 84Q ataxin-3, and *ATG5* knockdown similarly prevented compound-induced mutant ataxin-3 lowering. However, AN2 and GW5074 did not markedly alter p62 abundance, the LC3-II/LC3-I ratio, or GFP-LC3B puncta number or size, whereas CQ induced GFP-LC3B puncta accumulation as expected (Fig. 4). These results indicate that AN2 and GW5074 selectively enhanced the autophagic degradation of mAtxn3 without activating the global autophagy, likely by inducing mAtxn-3-LC3 proximity.

**Fig. 4.**
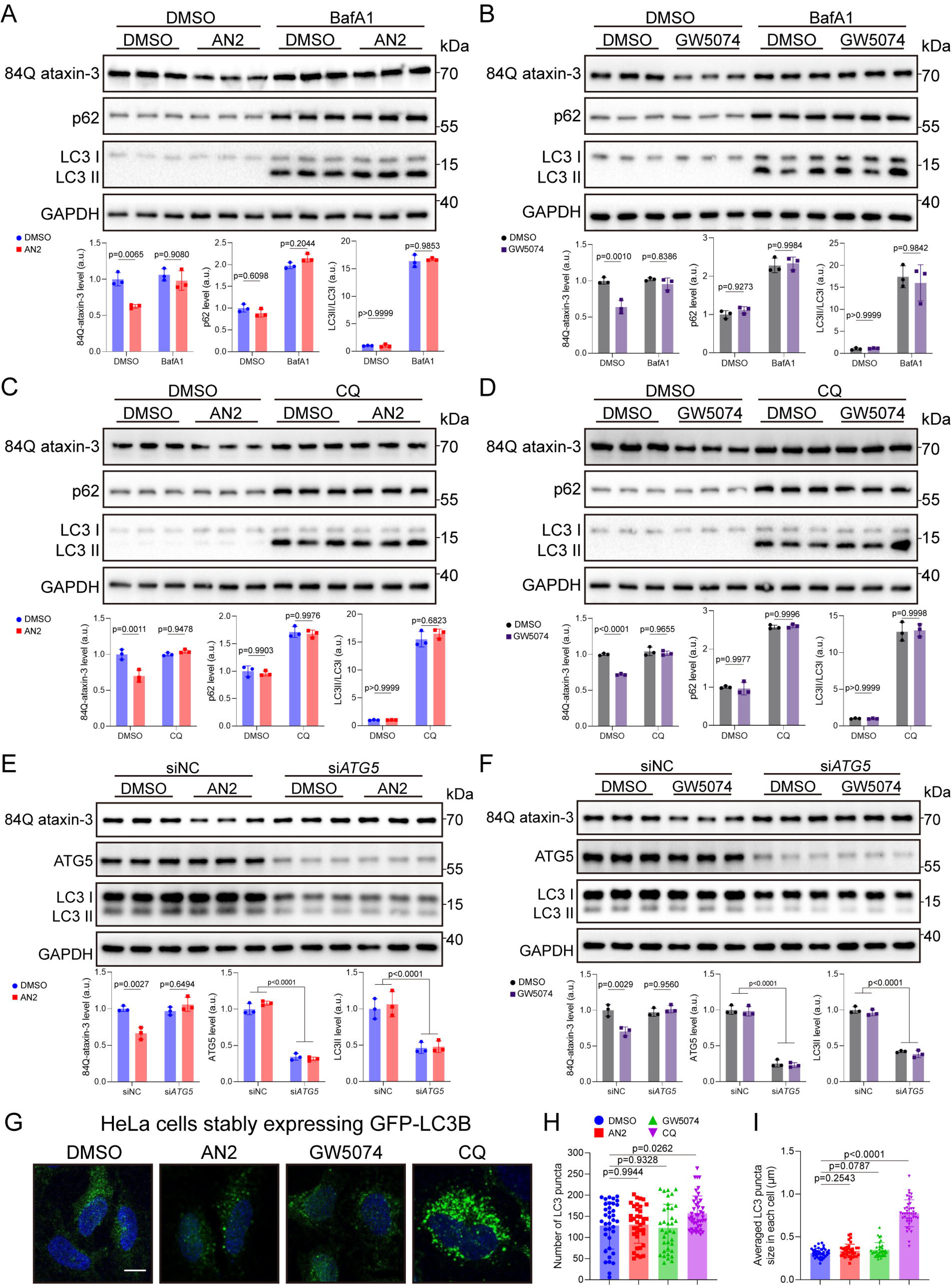
AN2 and GW5074 reduce mutant ataxin-3 through an autophagy-dependent mechanism without overt activation of bulk autophagy. **(A and B)** Representative immunoblots and quantification of 84Q ataxin-3, p62, and the LC3-II/LC3-I ratio in SH-SY5Y cells stably expressing ATXN3-84Q-FLAG. Cells were pretreated with or without BafA1 (100nM) for 12 hours and then treated with DMSO, AN2 (100nM) **(A)**, or GW5074 (100nM) **(B)** for 48 hours. Protein abundance was normalized to GAPDH where appropriate and then to the mean value of the no-BafA1 DMSO group. Data are shown as mean ± SD; n = 3. Statistical analysis was performed by two-way ANOVA followed by Sidak’s multiple-comparisons test comparing DMSO versus compound treatment within each BafA1 condition. For 84Q ataxin-3, the interaction *P* values between BafA1 treatment and compound treatment were P **=** 0.0238 for AN2 and P = 0.0050 for GW5074. **(C and D)** Representative immunoblots and quantification of 84Q ataxin-3, p62, and the LC3-II/LC3-I ratio in SH-SY5Y cells stably expressing ATXN3-84Q-FLAG. Cells were pretreated with or without CQ (40μM) for 12 hours and then treated with DMSO, AN2 (100nM) **(C)**, or GW5074 (100nM) **(D)** for 48 hours. Protein abundance was normalized to GAPDH where appropriate and then to the mean value of the no-CQ DMSO group. Data are shown as mean ± SD; n = 3. Statistical analysis was performed by two-way ANOVA followed by Sidak’s multiple-comparisons test comparing DMSO versus compound treatment within each CQ condition. For 84Q ataxin-3, the interaction *P* values between CQ treatment and compound treatment were P = 0.0007 for AN2 and P = 0.0002 for GW5074. **(E and F)** Representative immunoblots and quantification of 84Q ataxin-3, ATG5, and LC3-II in SH-SY5Y cells stably expressing ATXN3-84Q-FLAG after transfection with control siRNA (siNC) or si*ATG5*, followed by treatment with DMSO, AN2 (100nM) **(E)**, or GW5074 (100nM) **(F)** for 48 hours. Protein abundance was normalized to GAPDH and then to the mean value of the siNC + DMSO group. Data are shown as mean ± SD; n = 3. Statistical analysis for 84Q ataxin-3 was performed by two-way ANOVA followed by Sidak’s multiple-comparisons test comparing DMSO versus compound treatment within each siRNA condition. The interaction *P* values between siRNA treatment and compound treatment were P = 0.0009 for AN2 and P = 0.0017 for GW5074. ATG5 and LC3-II quantification was used to confirm *ATG5* knockdown and the associated reduction in LC3-II. **(G)** Representative confocal images of HeLa cells stably expressing GFP-LC3B after treatment with DMSO, AN2 (100nM), GW5074 (100nM), or CQ (40μM) for 48 hours. CQ was used as a positive control for GFP-LC3 puncta accumulation. Green, GFP-LC3B; blue, DAPI. Scale bar, 10μm. **(H and I)** Quantification of GFP-LC3-positive puncta number **(H)** and average GFP-LC3 puncta size per cell **(I)** from **(G)**. Data are shown as mean ± SD; n = 36 to 43. GFP-LC3 puncta number was analyzed by one-way ANOVA followed by Dunnett’s multiple-comparisons test versus the DMSO group. Average GFP-LC3 puncta size was analyzed by Welch’s one-way ANOVA followed by Dunnett’s T3 multiple-comparisons test versus the DMSO group. *P* values are shown in the plots.

### AN2 and GW5074 lowered mAtxn3 in MJD84.2 SCA3 mice after central and intraperitoneal delivery

We next evaluated AN2 and GW5074 in vivo in the well-established MJD84.2 SCA3 transgenic mice, which express human mutant *ATXN3* and develop progressive motor and neuropathological abnormalities [20]. Transgenic (TG) mice were confirmed by genotyping and showed human mutant ataxin-3 expression, ataxin-3-positive nuclear inclusions in the deep cerebellar nuclei and pons, and motor deficits by 15 months of age (Supplementary Fig. 4, A to E). To test central nervous system target engagement, 8-month-old male TG mice received intracerebroventricular injections of DMSO, AN2 (25μM), or GW5074 (25μM) for 10 days. Both compounds reduced human mutant ataxin-3 in the cerebellum, brainstem, and cerebral cortex without detectably altering endogenous mouse ataxin-3, demonstrating in vivo target selectivity after central delivery (Supplementary Fig. 4, F to H).

We then tested peripheral intraperitoneal administration in 15-month-old male and female wild-type (WT) or TG mice treated with DMSO, AN2 (0.5mg/kg), or GW5074 (0.5mg/kg) once daily for 14 days. AN2 and GW5074 improved rotarod and balance beam measures in TG mice of both sexes, while showing no detectable behavioral effect in WT mice. Both compounds also reduced human mutant ataxin-3 across the cerebellum, brainstem, and cerebral cortex of male and female TG mice, with no detectable change in endogenous wild-type mouse ataxin-3, consistent with the in vitro observations (Supplementary Figs. 5 and 6).

**Fig. 5.**
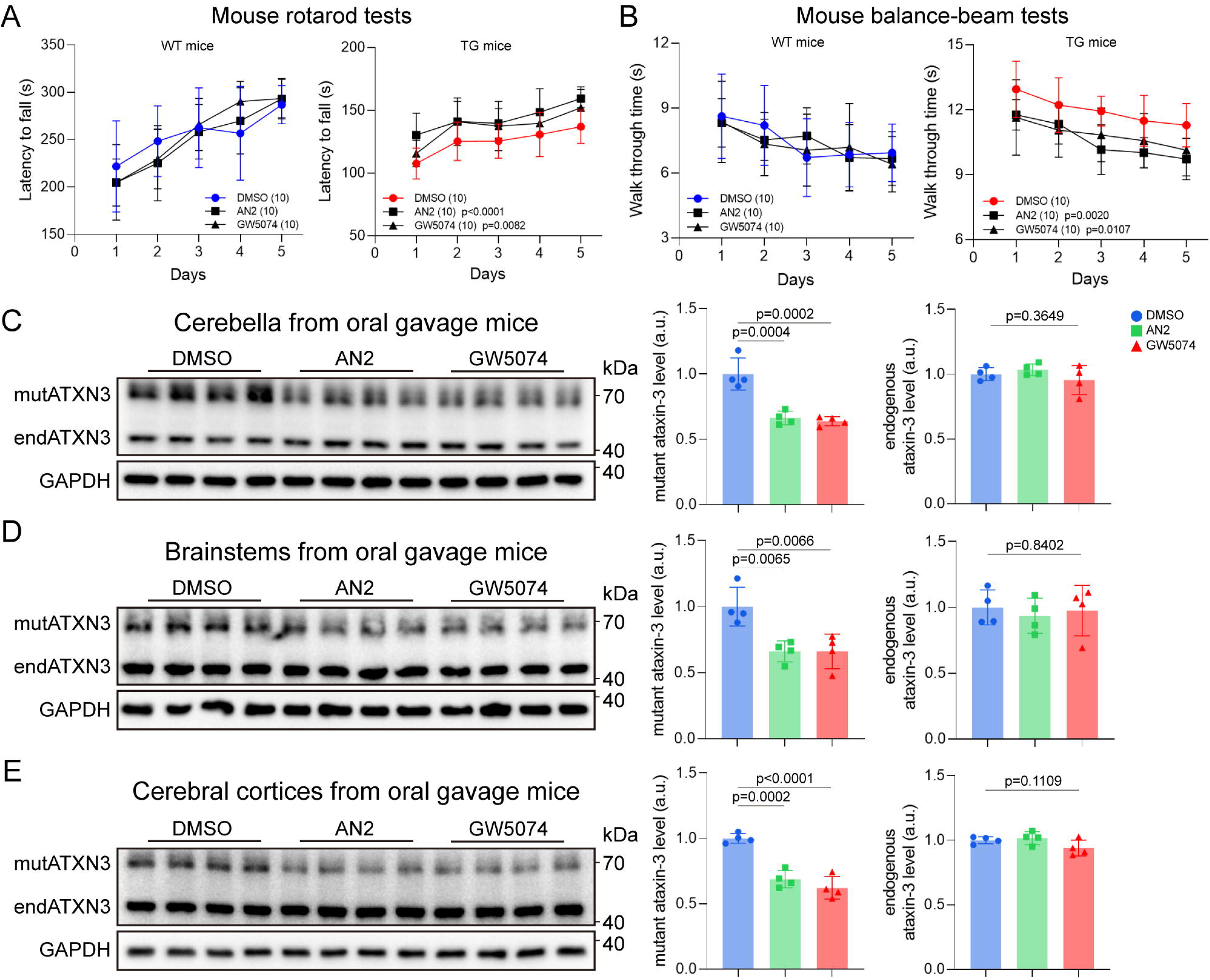
Oral AN2 and GW5074 improve motor phenotypes and reduce mutant ataxin-3 in male SCA3 transgenic mice. Fifteen-month-old male wild-type (WT) or SCA3 transgenic (TG) mice in the indicated experimental groups were orally treated with DMSO, AN2 (15mg/kg), or GW5074 (15mg/kg) for 28 days. **(A and B)** Rotarod performance **(A)** and balance-beam performance **(B)** of WT and TG mice after oral treatment. Rotarod performance is shown as latency to fall, and balance-beam performance is shown as traversal time. For each mouse, the value represents the mean of three trials. Data are shown as mean ± SD; n = 10. Statistical analysis was performed using two-way repeated-measures ANOVA with Dunnett-adjusted comparisons of the overall treatment effects versus DMSO. *P* values shown in the plots represent comparisons across all time points. The *P* values shown in the legends indicate the overall treatment effects compared with DMSO. **(C to E)** Representative immunoblots and quantification of mutant human ataxin-3 and endogenous mouse ataxin-3 in the cerebellum **(C)**, brainstem **(D)**, and cerebral cortex **(E)** of treated male TG mice. Protein abundance was normalized to GAPDH and then to the mean value of the DMSO group. Data are shown as mean ± SD; n = 4. Statistical analysis was performed by one-way ANOVA followed by Dunnett’s multiple-comparisons test versus the DMSO group. mutATXN3, mutant human ataxin-3; endATXN3, endogenous mouse ataxin-3.

**Fig. 6.**
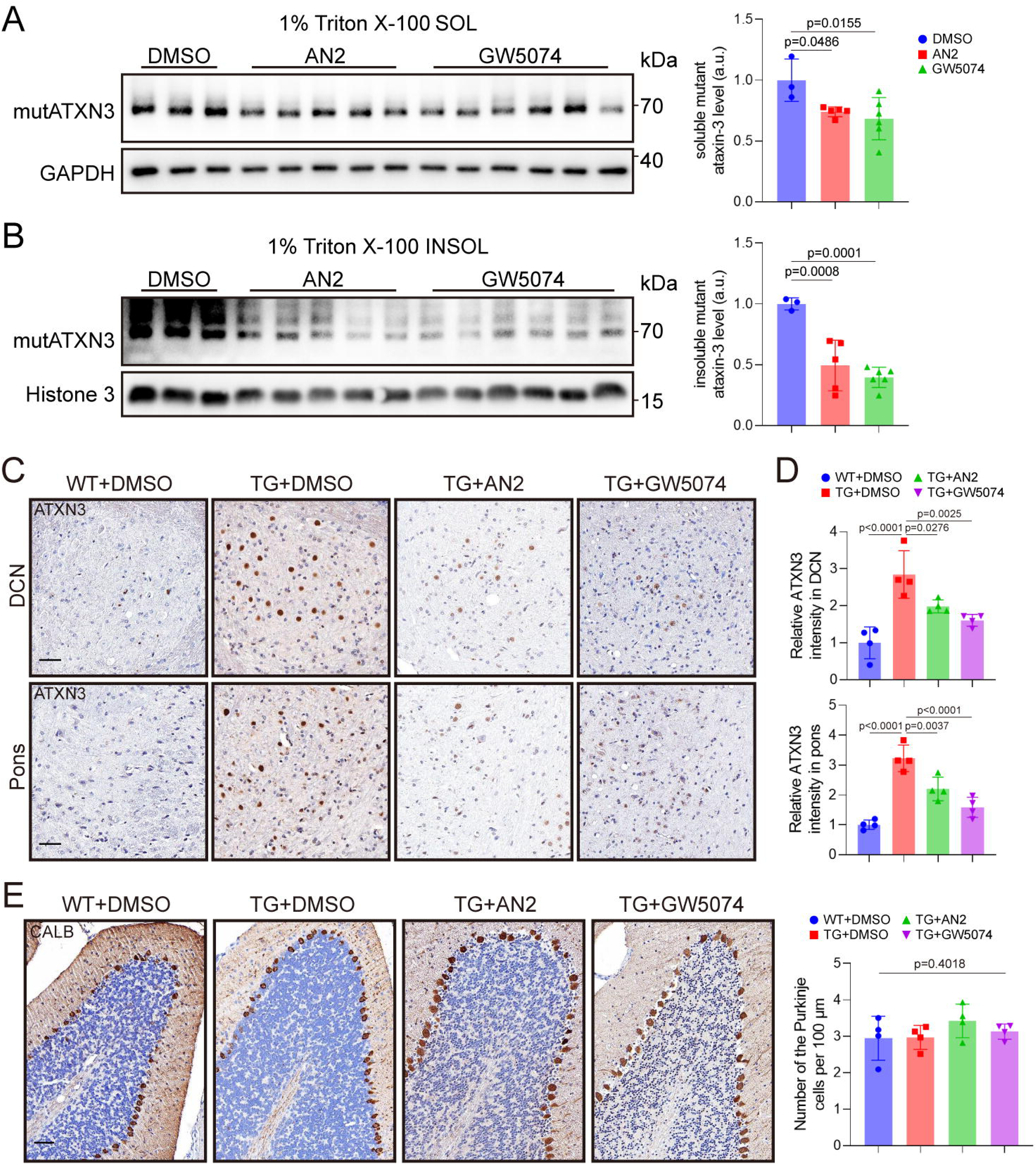
Oral AN2 and GW5074 preferentially reduce insoluble mutant ataxin-3 and ameliorate cerebellar pathology in SCA3 transgenic mice **(A and B)** Representative immunoblots and quantification of mutant human ataxin-3 in 1% Triton X-100-soluble **(A)** and 1% Triton X-100-insoluble **(B)** cerebellar fractions from 15-month-old SCA3 transgenic (TG) mice orally treated with DMSO, AN2 (15mg/kg), or GW5074 (15mg/kg) for 28 days. Protein abundance was normalized to GAPDH for soluble fractions and to Histone 3 for insoluble fractions, and then to the mean value of the DMSO group. Data are shown as mean ± SD; n = 3 to 6. Statistical analysis was performed by one-way ANOVA followed by Dunnett’s multiple-comparisons test versus the DMSO group. SOL, Triton X-100-soluble fraction; INSOL, Triton X-100-insoluble fraction; mutATXN3, mutant human ataxin-3. **(C and D)** Representative ataxin-3 immunohistochemistry images **(C)** and quantification of relative ataxin-3 immunoreactivity **(D)** in the deep cerebellar nuclei (DCN) and pons of 15-month-old wild-type (WT) and TG mice after 28 days of oral treatment. Quantification was normalized to the mean value of the WT + DMSO group. **(E)** Representative calbindin immunohistochemistry images and quantification of Purkinje cell number in the cerebellum after oral treatment. Scale bars, 50μm. For **(D)** and **(E)**, data are shown as mean ± SD; n = 4. Statistical analysis was performed by one-way ANOVA followed by Dunnett’s multiple-comparisons test versus the TG + DMSO group. CALB, calbindin.

### Oral AN2 and GW5074 administrations were tolerated and therapeutically active in SCA3 transgenic mice

Because oral administration is more clinically acceptable and patient-friendly than intracerebral or intraperitoneal delivery, we next evaluated oral AN2 and GW5074. After a single 15mg/kg oral gavage dose in WT mice, both compounds were detected in plasma and cerebellum, with cerebellar concentrations of approximately 20 to 150nM for AN2 and 20 to 80nM for GW5074, overlapping the effective in vitro concentration range for mAtxn3 reduction. GW5074 declined more rapidly than AN2 in cerebellar tissue; therefore, subsequent oral studies used once-daily AN2 and three-times-daily GW5074. Under these dosing regimens, 28-day oral treatment in WT mice did not cause overt abnormalities in body weight, serum liver- or kidney-function-related biochemical indices, liver or kidney weight-to-body weight ratios, or liver and kidney histology, suggesting relatively good tolerability of these compounds (Supplementary Fig. 7).

We then treated 15-month-old SCA3 TG mice by oral gavage for 28 days. In both male and female TG mice, AN2 and GW5074 improved rotarod and balance beam measures, whereas the same treatments did not detectably alter rotarod or balance beam performance in WT mice. Both compounds also reduced human mutant ataxin-3 in the cerebellum, brainstem, and cerebral cortex, with little detectable effect on endogenous wild-type mouse ataxin-3 (Fig. 5; Supplementary Fig. 8).

Cerebellar biochemical fractionation showed that oral AN2 and GW5074 reduced mutant ataxin-3 in both Triton X-100-soluble and Triton X-100-insoluble fractions, with a stronger effect in the insoluble fraction, consistent with the cellular aggregation-associated protein results (Fig. 6, A and B). Oral treatment also reduced ataxin-3 immunoreactivity in the deep cerebellar nuclei and pons (Fig. 6, C to D). During the same treatment period, AN2 and GW5074 did not further increase GFAP or IBA1 signals in the deep cerebellar nuclei or pons and did not significantly alter selected off-target proteins in cerebellar lysates, including the wild-type versions of the polyQ proteins including ataxin-2, Thap11, and Htt, and the house-keeping proteins including β-actin, β-tubulin, and GAPDH (Supplementary Fig. 9). Together, these findings indicate that oral AN2 and GW5074 achieved brain exposure, selectively lowered mAtxn3, preferentially reduced insoluble mAtxn-3 species, and improved motor and neuropathological phenotypes without overt toxicity under the 28-day dosing regimens tested.

### SCA3 iPSC-derived neurons confirmed allele-selective mAtxin-3 lowering

To further validate the therapeutic potential of these compounds, we tested the activity of AN2 and GW5074 in a human neuronal model. We generated iPSCs from peripheral blood mononuclear cells obtained from one male patient with SCA3 and one age- and sex-matched healthy control. The SCA3 iPSC line carried *ATXN3* alleles with 15 and 78 CAG repeats, whereas the control line carried 28 and 29 repeats. Both iPSC lines showed typical colony morphology, alkaline phosphatase activity, normal karyotypes, absence of mycoplasma contamination, and expression of pluripotency markers. NGN2-based induction generated TUJ1- and MAP2-positive neurons by differentiation day 14, and neurons differentiated from SCA3 iPSCs expressed both wild-type and mutant ataxin-3 (Supplementary Fig. 10).

During early neuronal differentiation, AN2 and GW5074 did not detectably alter wtAtaxin-3 in healthy control neurons but allele-selectively reduced mAtxn-3 in SCA3 neurons (Fig. 7A; Supplementary Fig. 11, A and B). CQ pretreatment blocked this mAtxn-3-lowering effect, whereas p62 abundance and the LC3-II/LC3-I ratio were not markedly altered within matched CQ conditions (Supplementary Fig. 11, C and D). AN2 and GW5074 also did not detectably change selected wild-type polyQ proteins, including ataxin-2, THAP11, and HTT, or loading-control proteins in SCA3 neurons (Fig. 7, B and C). Together, these findings support allele-selective and autophagy-dependent mutant ataxin-3 lowering in patient-derived neurons.

**Fig. 7.**
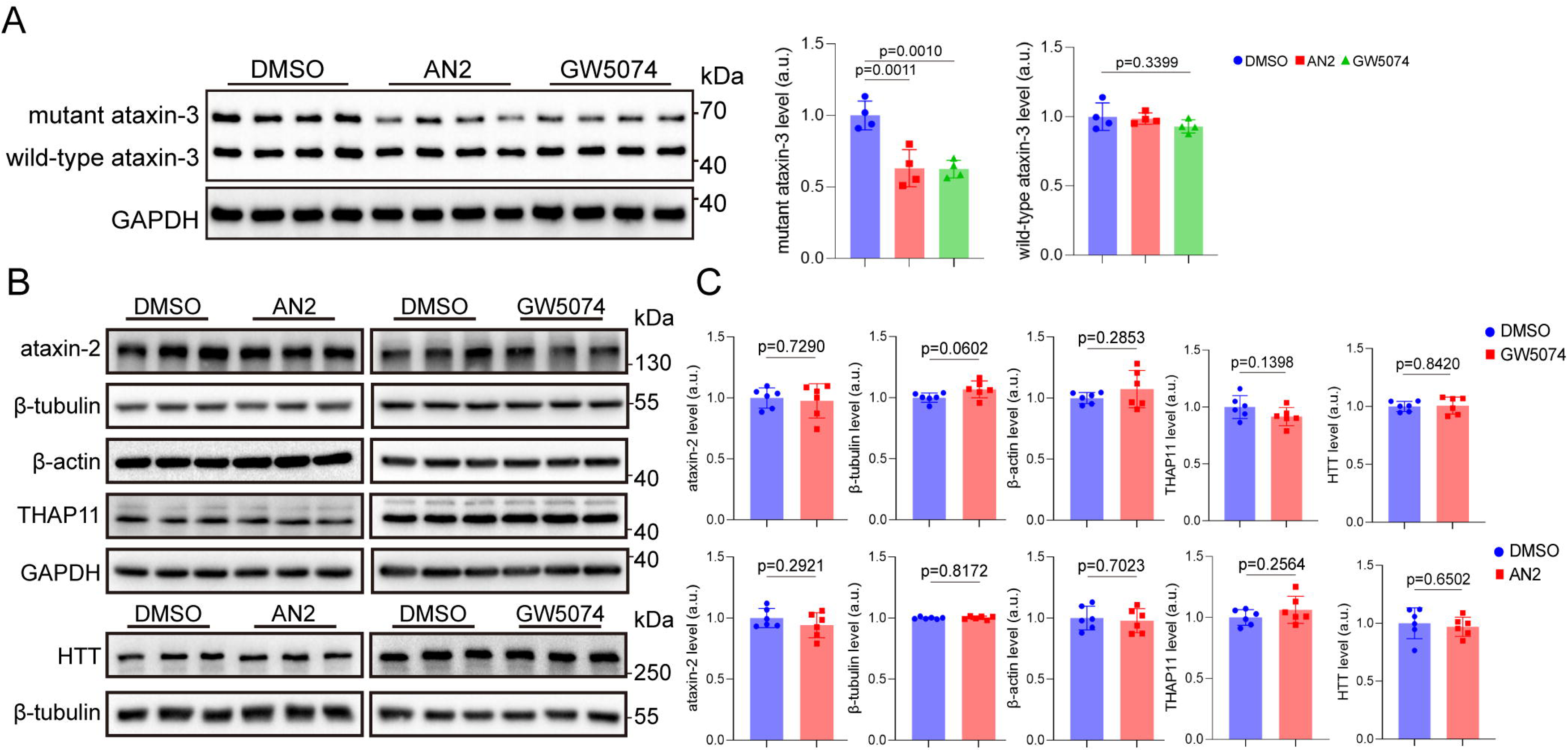
AN2 and GW5074 reduce mutant ataxin-3 but do not broadly alter selected nontarget proteins in SCA3 iPSC-derived neurons. **(A)** Representative immunoblot and quantification of ataxin-3 in cortical neurons differentiated from SCA3 patient-derived iPSCs after treatment with DMSO, AN2 (100nM), or GW5074 (100nM). Protein abundance was normalized to GAPDH and then to the mean value of the DMSO group. Data are shown as mean ± SD; n = 4. Statistical analysis was performed by one-way ANOVA followed by Dunnett’s multiple-comparisons test versus the DMSO group. **(B and C)** Representative immunoblots and quantification of selected nontarget proteins in cortical neurons differentiated from SCA3-iPSCs after treatment with DMSO, AN2 (100nM), or GW5074 (100nM) until differentiation day 5. **(B)** Immunoblots showing ataxin-2, THAP11, β-actin, β-tubulin, and GAPDH, as well as HTT and β-tubulin. (**C**) Quantification shows the levels of ataxin-2, THAP11, HTT, and loading-control proteins under the indicated treatment conditions. Target protein abundance was normalized to GAPDH or β-tubulin and then to the mean value of the corresponding DMSO group. Data are shown as mean ± SD; n = 6. Statistical analysis was performed by two-tailed Student’s *t* test.

### PolyQ length analyses supported expanded polyQ-dependent lowering of ataxin-3

Finally, we examined whether AN2 and GW5074 activity was related to *ATXN3* CAG-repeat length. In 1303 Chinese patients with SCA3, expanded *ATXN3* alleles ranged from 48 to 83 CAG repeats, whereas normal alleles ranged from 11 to 36 repeats, with 90.02% of normal alleles containing 27 or fewer repeats (Fig. 8, A and B). We then expressed ATXN3 constructs encoding 74Q, 57Q, 46Q, 40Q, or 27Q ataxin-3 in HEK293T cells and treated the cells with AN2 or GW5074 at 50, 100, or 200nM for 48 hours. Both compounds reduced 74Q, 57Q, 46Q, and 40Q ataxin-3 but did not detectably reduce 27Q ataxin-3, supporting polyQ length-dependent activity of these compounds with relative sparing of an ataxin-3 polyQ length range that covered most (>90%) normal *ATXN3* alleles in this Chinese SCA3 cohort (Fig. 8, C to G).

**Fig. 8.**
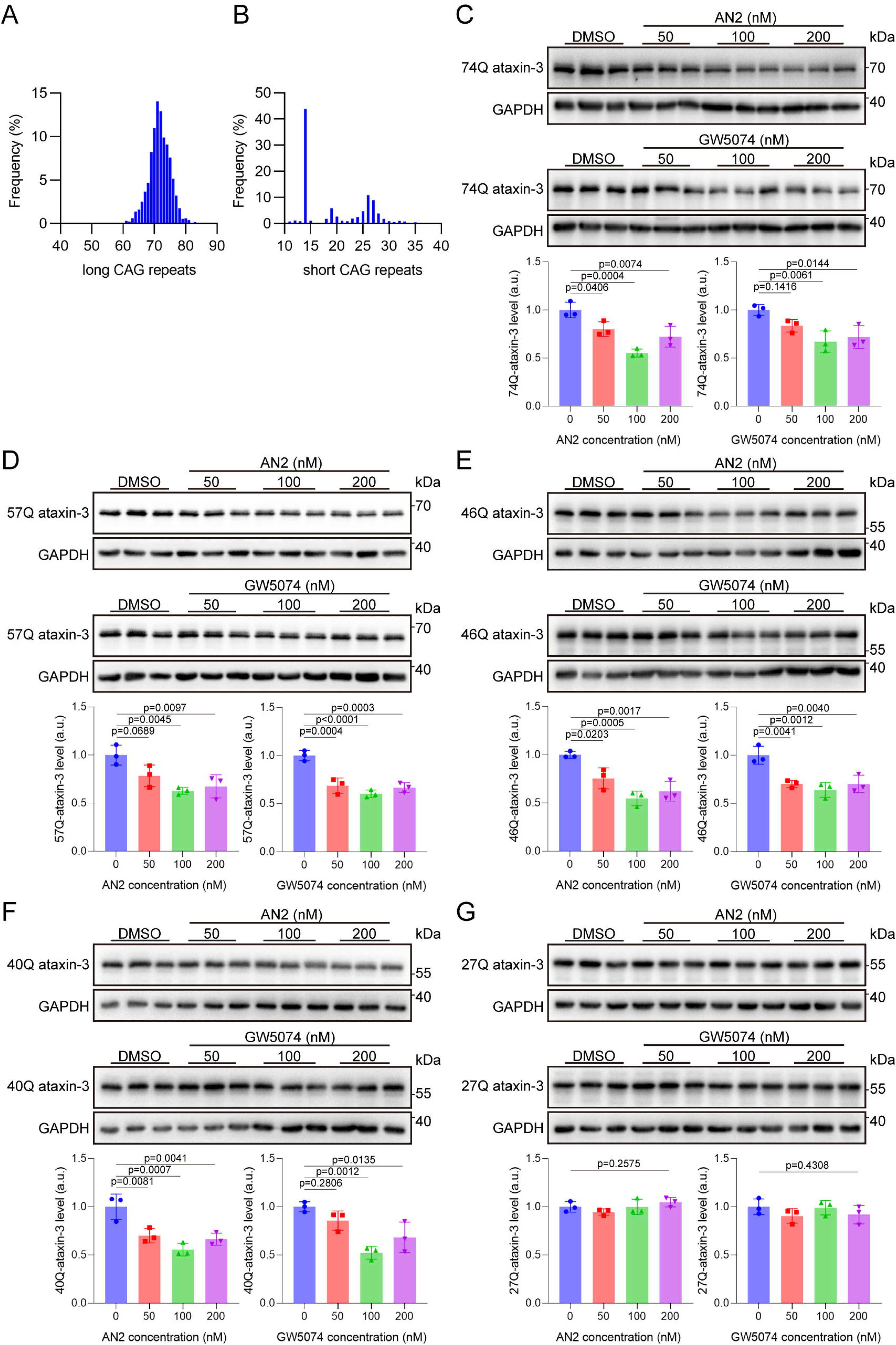
*ATXN3* CAG-repeat distribution in a Chinese SCA3 cohort and polyQ length-dependent lowering of ataxin-3 by AN2 and GW5074. **(A and B)** Frequency distributions of CAG-repeat lengths in the expanded **(A)** and normal **(B)** *ATXN3* alleles in the Chinese SCA3 cohort. **(C to G)** Representative immunoblots and quantification of ataxin-3 with different polyQ lengths in HEK293T cells after treatment with DMSO or the indicated concentrations of AN2 or GW5074 for 48 hours. Panels show 74Q ataxin-3 **(C)**, 57Q ataxin-3 **(D)**, 46Q ataxin-3 **(E)**, 40Q ataxin-3 **(F),** and 27Q ataxin-3 **(G)**. Protein abundance was normalized to GAPDH and then to the mean value of the DMSO group. Data are shown as mean ± SD; n = 3. Statistical analysis was performed by one-way ANOVA followed by Dunnett’s multiple-comparisons test versus the DMSO group. *P* values are shown in the plots.

## Discussion

This study provides preclinical proof of concept that mutant ataxin-3 can be selectively lowered through an autophagosome-tethering strategy in SCA3. Previous therapeutic studies in SCA3 have mainly pursued *ATXN3* transcript suppression or enhancement of intracellular protein clearance, including antisense oligonucleotides, RNA interference, oligonucleotide-based approaches, and autophagy induction [10, 21–25]. These studies establish mutant ataxin-3 lowering as a disease-relevant therapeutic direction, but they also highlight unresolved translational issues, including central nervous system delivery, allele selectivity, durability, and long-term safety. Different from those gene therapy approaches, AN2 and GW5074 act at the protein-clearance level. Our data show that these compounds selectively lower polyQ-expanded ataxin-3 across overexpression cell models, MJD84.2 transgenic mice by oral administrations, and SCA3 patient-derived cortical neurons, providing potential entry points of developing orally available drugs for SCA3. Mechanistically, the study also suggest the mAtxn3 protein with the expanded polyQ as an important driver of the disease, independent from pathogenic mechanisms at the DNA or mRNA level, such as somatic expansion and RNA phase-separation [9, 18, 19].

A key implication of this work is that mutant ataxin-3 lowering can be achieved without broad stimulation of bulk autophagy. Pharmacological autophagy induction, including temsirolimus treatment, has been shown to reduce mutant ataxin-3 and improve phenotypes in SCA3 models, supporting the therapeutic relevance of the autophagy-lysosome pathway [10]. However, global enhancement of autophagy has recognized limitations in neurodegenerative disease therapy and may affect substrates beyond the intended disease protein [26]. In the present study, BafA1, CQ, and *ATG5* knockdown blocked compound-induced mutant ataxin-3 lowering, whereas p62 abundance, the LC3-II/LC3-I ratio, and GFP-LC3B puncta were not markedly altered at active compound concentrations. Because LC3-based readouts require cautious interpretation and do not alone define autophagic flux, these data should not be overinterpreted as a complete flux analysis [27]. Nevertheless, together with TurboID proximity labeling and colocalization imaging, they support a model in which AN2 and GW5074 enhance the spatial association between LC3 and polyQ-expanded ataxin-3 and use intact autophagy machinery for selective substrate degradation rather than acting as general autophagy activators.

The preferential reduction of insoluble and aggregation-associated mutant ataxin-3 is another important feature of this strategy. SCA3 pathology is characterized by misfolding, nuclear accumulation, and inclusion formation of expanded ataxin-3 in vulnerable neuronal populations [28, 29]. In our study, AN2 and GW5074 reduced mutant ataxin-3 in both soluble and insoluble fractions, with a stronger effect in the insoluble fraction in cells and a similar pattern in cerebellar tissue after oral dosing. The reduction in SDS-resistant species and aggregate-positive cells further suggests activity against biochemical and morphologic features of pathological ataxin-3 accumulation. These data do not identify which ataxin-3 conformer is most toxic or define the amount of lowering required for durable disease modification. However, the convergence of reduced aggregation-associated ataxin-3 with improved cell viability, ATP content, motor performance and ataxin-3 immunopathology supports the biological relevance of lowering this disease-associated protein pool. Notably, the observation that the compounds preferentially lowered aggregated mAtxn-3 is somewhat different from the observation on mHtt, of which the soluble form was degraded more by these compounds [9], possibility reflecting the difference between the aggregation properties of mHtt and mAtxn3 that may worth future investigation.

The in vivo findings strengthen the translational argument for allele-selective mAtxn-3 degradation. MJD84.2 mice express human mutant *ATXN3* and have been widely used for molecular and therapeutic studies in SCA3 [20]. In this model, AN2 and GW5074 reduced human mutant ataxin-3 after intracerebroventricular injection, intraperitoneal injection, and oral gavage, indicating that target engagement was not confined to a single delivery route. Oral treatment is particularly relevant because both compounds achieved nanomolar cerebellar exposure overlapping the effective cellular concentration range and improved behavioral and neuropathological readouts after 28 days. The absence of overt body weight loss, liver or kidney abnormalities, glial activation in the analyzed regions, or changes in selected nontarget proteins provides initial tolerability support. These experiments are not a substitute for formal toxicology, but they establish a practical basis for medicinal chemistry optimization, longer dosing studies, and more comprehensive safety profiling.

The patient-derived neuron and CAG-repeat analyses add a human-facing dimension to this preclinical study. Previous studies of ATTEC compounds did not evaluate the portion of polyQ patient populations that are potentially suitable for treatment by these compounds, which is critical for clinical translation of these compounds. In SCA3 iPSC-derived cortical neurons carrying 15/78 *ATXN3* CAG repeats, AN2 and GW5074 lowered mutant ataxin-3 while sparing the normal allele. This is notable because allele-selective lowering has also been a major objective of ASO-based SCA3 studies in patient-derived neuronal systems [25]. The cohort analysis further showed a clear separation between most normal alleles and expanded pathogenic alleles in 1303 Chinese patients with SCA3: 90.02% of normal alleles contained 27 or fewer CAG repeats, whereas expanded alleles ranged from 48 to 83 repeats. Consistently, AN2 and GW5074 reduced 40Q to 74Q ataxin-3 but did not detectably reduce 27Q ataxin-3. These findings support polyQ length-dependent activity with relative sparing of ataxin-3 lengths common among normal alleles in this cohort. Defining the 28Q to 39Q interval will be important because this range may determine the practical boundary between normal, intermediate, and compound-sensitive polyQ lengths.

This study has several limitations. The 28-day oral treatment window in mice was short relative to a lifelong neurodegenerative disease. Off-target assessment was limited to selected polyQ and housekeeping proteins; unbiased proteomic and transcriptomic analyses, together with long-term dosing and safety studies, will be required before clinical translation.

In summary, this study establishes allele-selective autophagic degradation of mAtxn-3 by small molecule compounds as a promising strategy for SCA3 treatment. The two proof-of-concept compounds provide useful entry points of oral drug development and chemical tools for mechanistic study of mAtxn-3 protein with the expanded polyQ.

## Materials and Methods

### Study design

This preclinical study evaluated whether the autophagosome-tethering compounds AN2 and GW5074 reduce mutant ataxin-3 and ameliorate SCA3-related phenotypes. Cellular studies used HEK293T transient expression models, SH-SY5Y stable expression models, HeLa imaging models, and SCA3 patient-derived iPSC cortical neurons. In vivo studies used MJD84.2 transgenic mice and wild-type littermates to assess target engagement, motor behavior, oral brain exposure, and tolerability. Sample sizes, replicate numbers, and statistical tests are reported in the figure legends.

### Plasmids and reagents

FLAG-tagged human ATXN3-20Q and ATXN3-84Q expression plasmids were used to generate cellular SCA3 models, with empty vector as a control where indicated. Additional pCDH-CMV-ATXN3-FLAG constructs encoding 74Q, 57Q, 46Q, 40Q, or 27Q ataxin-3 were used for polyQ-length response experiments. For stable SH-SY5Y cell lines, pCDH-CMV-ATXN3-20Q-FLAG and pCDH-CMV-ATXN3-84Q-FLAG lentiviral plasmids were packaged with psPAX2 (Addgene, 12260) and pMD2.G (Addgene, 12259). Additional plasmids included EGFP-ATXN3-20Q, EGFP-ATXN3-84Q, ATXN3-20Q-HA-TurboID, ATXN3-84Q-HA-TurboID, RFP-LC3, and pCMV-GFP-LC3B. Plasmids were amplified in DH5α competent cells (Tsingke, DLC114) and purified using an endotoxin-free plasmid extraction kit (Simgen, 1006050). AN2 (TargetMol, T5070) and GW5074 (dcChemicals, DC8810) were dissolved in DMSO (Solarbio, D8371) to prepare concentrated stock solutions, which were further diluted to intermediate working stocks before use. Chloroquine (CQ; Selleck, S6999) and bafilomycin A1 (BafA1; Selleck, S1413) were used for autophagy-inhibition experiments. Antibody sources, catalog numbers, and dilutions are listed in Supplementary Table 1.

### Cell culture, transfection, and stable cell lines

HEK293T or HeLa cells were maintained in high-glucose DMEM (Servicebio, G4523) supplemented with 10% fetal bovine serum (FBS; Procell, 164210) and 1% penicillin-streptomycin (Procell, PB180120). SH-SY5Y cells were maintained in MEM/F12 medium (Procell, PM151220) supplemented with 15% FBS and 1% penicillin-streptomycin. Cells were cultured at 37 in 5% CO_2_. For transient expression experiments, HEK293T or HeLa cells were transfected at 50 to 60% confluence using Lipofectamine 3000 (Invitrogen, L3000075) according to the manufacturer’s protocol. Cells were used for subsequent experiments 24 to 48 hours after transfection, as indicated.

SH-SY5Y cells stably expressing ATXN3-20Q-FLAG or ATXN3-84Q-FLAG were generated by lentiviral transduction. Viral supernatants were produced in HEK293T cells, collected 48 hours after transfection, centrifuged at 3000rpm for 10min, filtered through a 0.45μm filter, and used to infect SH-SY5Y cells in the presence of 8μg/ml polybrene (Sigma-Aldrich, H9268). Cells were selected with 2μg/ml puromycin (Selleck, S7417) for 1 to 2 weeks, and stable expression was confirmed by immunoblotting. HeLa cells stably expressing GFP-LC3B were generated by lentiviral transduction and puromycin selection using the same procedure.

### Compound treatment, autophagy inhibition, and *ATG5* knockdown

For FLAG-tagged ATXN3 concentration-response experiments, cells were treated with DMSO, AN2, or GW5074 at 30, 50, 100, 200, or 300nM for 48 hours. For polyQ-length response experiments, cells were treated with 50, 100, or 200nM compound for 48 hours. Validation experiments used 100nM AN2 or GW5074 unless otherwise indicated. For pharmacological autophagy inhibition, SH-SY5Y cells stably expressing ATXN3-84Q-FLAG were pretreated with BafA1 (100nM) or CQ (40μM) for 12 hours, followed by treatment with DMSO, AN2, or GW5074 treatment for 48 hours in the continued presence of the inhibitor. For *ATG5* knockdown, cells were transfected with control siRNA or si*ATG5* using Lipofectamine RNAiMAX (Invitrogen, 13778150); the si*ATG5* target sequence was GCCUGUAUGUACUGCUUUA. Forty-eight hours after siRNA transfection, cells were treated with DMSO, AN2, or GW5074 for an additional 48 hours before harvest.

For biochemical aggregation experiments, HEK293T cells were transiently transfected with EGFP-ATXN3-20Q or EGFP-ATXN3-84Q plasmids. To assess the formation of SDS-resistant ataxin-3 species, cells were harvested on days 3, 5, and 7 after transfection. To test compound effects on SDS-resistant species, EGFP-ATXN3-expressing cells were treated continuously with DMSO, AN2 (100nM), or GW5074 (100nM) for 7 days after transfection, followed by biochemical fractionation and western blotting.

### Protein extraction and western blotting

Cells or dissected brain regions were lysed in freshly prepared 2% SDS lysis buffer supplemented with protease inhibitor cocktail (Selleck, B14001) and phosphatase inhibitor cocktails A and B (Selleck, B15001) at a volume ratio of 100:1:1:1. Lysates were homogenized with 3-mm grinding beads using a low-temperature grinder and centrifuged at 12,000g for 20min at 4 . Supernatants were collected, and protein concentrations were determined using a BCA protein assay kit (Thermo Scientific, 23225). Samples were mixed with 6× loading buffer and heated at 98 for 10min. Proteins were separated by SDS-PAGE and transferred onto 0.2μm PVDF membranes (Millipore). Membranes were blocked with 5% nonfat milk in TBST, incubated with primary antibodies overnight at 4, and then incubated with HRP-conjugated secondary antibodies. Bands were detected using ECL substrates (ShareBio, SB-WB012 or SB-WB011) and imaged with a Bio-Rad chemiluminescence system. Band intensities were quantified using ImageJ and normalized to the indicated loading control and then to the indicated control group.

### Triton X-100/SDS fractionation

Triton X-100/SDS sequential fractionation was used to separate soluble and insoluble ataxin-3 species, with modifications from a published ATXN3 biochemical fractionation protocol [30]. Cells were lysed in 500μl of freshly prepared 1% Triton X-100 lysis buffer supplemented with protease and phosphatase inhibitors, incubated on ice for 30min, and centrifuged at 15,000g for 20min at 4 . Supernatants were collected as Triton X-100-soluble fractions and sonicated on ice for 4min at 15% power with 3-s on/3-s off pulses. Pellets were washed twice with 1ml of 1% Triton X-100 lysis buffer and centrifuged at 15,000rpm for 5min at 4 after each wash. Washed pellets were resuspended in 250μl of freshly prepared 4% SDS lysis buffer, sonicated using the same settings, heated at 98 for 5 min, and collected as Triton X-100-insoluble fractions. Cerebellar fractionation followed the same sequential extraction principle, using 500μl of 1% Triton X-100 buffer and 300μl of 4% SDS buffer per sample. For SDS-resistant ataxin-3 analysis, fractions from EGFP-ATXN3-expressing HEK293T cells were analyzed by long-exposure western blotting to detect high-molecular-weight SDS-resistant ataxin-3 signals. Protein concentrations were determined by BCA assay, and equal amounts of protein were loaded for western blotting within each comparison.

### Cell viability, ATP, aggregate imaging, and GFP-LC3 analysis

Cell viability was measured using enhanced CCK-8 reagent (Elabscience, E-CK-A362). Cells were seeded in 96-well plates, treated with DMSO or compounds for 48 hours, incubated with CCK-8 reagent in complete medium, and read at 450nm using an Agilent BioTek Synergy multimode microplate reader. Cellular ATP content was measured using an enhanced ATP assay kit (Beyotime, S0027). Cells were lysed on ice, lysates were centrifuged at 12,000g for 5min at 4, and luminescence was measured after mixing samples or standards with ATP detection working solution. ATP values were normalized to total protein concentration.

For aggregate imaging, HeLa cells expressing EGFP-Ataxin3Q28 (Addgene, 22122) or EGFP-Ataxin3Q84 (Addgene, 22123) were fixed after compound treatment, mounted with DAPI-containing antifade medium (Abcam, ab104139), and imaged by confocal microscopy. Aggregate-positive cells were quantified as the percentage of EGFP-expressing cells containing visible aggregates. GFP-LC3B puncta were assessed in HeLa cells stably expressing GFP-LC3B after treatment with DMSO, AN2, GW5074, or CQ; puncta number and size were quantified using ImageJ.

### TurboID proximity labeling and LC3 colocalization

HEK293T cells expressing HA-TurboID, ATXN3-20Q-HA-TurboID, or ATXN3-84Q-HA-TurboID were treated with 100μM biotin and DMSO, AN2, or GW5074 for 4 hours. Cells were lysed in inhibitor-supplemented RIPA buffer (Beyotime, P0013B), sonicated, and centrifuged. Input lysates were retained, and biotinylated proteins were enriched with streptavidin magnetic agarose beads (AlpaLifeBio, KTSM-K2405), washed, eluted with loading buffer, and analyzed by western blotting.

For colocalization analysis, HeLa cells were cotransfected with RFP-LC3 and EGFP-Ataxin3Q28 or EGFP-Ataxin3Q84, treated with compounds for 4 hours, fixed, and imaged by confocal microscopy. Colocalized puncta were calculated as the percentage of EGFP- and RFP-double-positive puncta among total EGFP-positive puncta.

### Animals and compound administration

All animal experiments were approved by the Animal Welfare and Ethics Committee of the Department of Laboratory Animals of Central South University and the Animal Ethics Committee of Xiangya Hospital, Central South University (approval numbers CSU-2023-0001 and 2024020019). MJD84.2 SCA3 transgenic (TG) mice, originally described as a human mutant *ATXN3* transgenic model, and wild-type (WT) littermates were used for in vivo studies [20]. Mice were maintained by crossing heterozygous TG mice with C57BL/6J WT mice and housed under specific pathogen-free conditions with a 12-hour light/dark cycle, standard chow, and water ad libitum. Genotypes were determined by PCR using tail DNA and primers detecting the human *ATXN3* transgene, as previously described [20]. Age- and sex-matched mice were used for the indicated experiments and assigned to treatment groups before dosing.

For intracerebroventricular (ICV) administration, 8-month-old male TG mice were implanted with a guide cannula targeting the lateral ventricle, based on a previously reported ICV delivery protocol [9], and treated once daily for 10 days with vehicle, AN2, or GW5074 (25μM in ACSF containing 0.125% DMSO; 2μl per injection at 0.25μl/min). For intraperitoneal administration, 15-month-old male and female WT or TG mice received vehicle, AN2, or GW5074 (0.5mg/kg) once daily for 14 days. For oral administration, AN2 and GW5074 were formulated in 4% DMSO, 30% PEG300, 5% Tween-80, and 61% water and administered by gavage at 15mg/kg. Based on cerebellar exposure profiles after a single oral dose, AN2 was dosed once daily and GW5074 three times daily in subsequent 28-day oral studies. For oral efficacy studies, 15-month-old male and female WT or TG mice were treated for 28 days and analyzed by sex where indicated; 15-month-old WT mice were used for the 28-day oral safety assessment.

### Measurement of compound concentrations

Plasma and cerebellar concentrations were measured after a single oral dose of 15mg/kg. Blood was collected by cardiac puncture, allowed to stand, and centrifuged at 3000rpm for 20min at room temperature to obtain plasma. Mice were perfused with PBS before cerebellar dissection. Plasma was extracted with methanol, and cerebellar tissues were homogenized in methanol with grinding beads. Supernatants were analyzed by liquid chromatography-mass spectrometry, and concentrations were calculated using standard curves generated with authentic AN2 or GW5074 standards.

### Behavioral testing

Motor performance was assessed using rotarod, and balance beam tests. Mice were habituated to the testing room and trained for 3 consecutive days before formal testing. In the rotarod test, mice were tested at 40 rpm with a maximum trial duration of 5 min. In the balance beam test, mice traversed a 0.5-cm-wide beam toward a dark box. For both tests, each mouse was tested in three trials separated by at least 30 min, and the mean value was used for analysis.

### Histology and immunostaining analysis

Mouse brains were fixed in 4% paraformaldehyde, embedded in paraffin, and sectioned at 5μm. Liver and kidney sections were stained with hematoxylin and eosin and imaged under bright-field microscopy. For brain immunohistochemistry, sections were subjected to antigen retrieval in Tris-EDTA buffer (pH 9.0; Servicebio, G1203), blocked, incubated with antibodies against ataxin-3 or calbindin, and developed using DAB substrate (Servicebio, G1212). DAB-stained sections were imaged under bright-field microscopy. For immunofluorescence, sections were incubated with antibodies against GFAP and IBA1, followed by fluorophore-conjugated secondary antibodies and DAPI counterstaining. Fluorescence images of the deep cerebellar nuclei and pons were acquired using a Zeiss fluorescence microscope.

### Human participants, iPSC generation, and neuronal induction

Peripheral blood samples were obtained from one male patient with SCA3 and one age- and sex-matched healthy control for iPSC generation. The SCA3 donor was 25 years old and carried *ATXN3* alleles with 15 and 78 CAG repeats; the control donor was 28 years old and carried 28 and 29 repeats. *ATXN3* CAG-repeat data from 1303 Chinese patients with SCA3 were used for cohort-level analysis. Written informed consent was obtained from all participants, and the study was approved by the Medical Ethics Committee of Xiangya Hospital, Central South University (202310206, 2024010125, and 2025030338). Peripheral blood mononuclear cells were reprogrammed using the CytoTune 2.0 Sendai virus kit (Thermo Scientific, A16517) according to the manufacturer’s protocol. iPSC colonies were selected, expanded, and characterized by morphology, alkaline phosphatase staining, mycoplasma testing, karyotyping, pluripotency marker staining, and *ATXN3* repeat analysis. Cortical neurons were generated using an NGN2-based induction protocol modified from published methods [31, 32]. For compound experiments, neurons were treated during early differentiation with DMSO, AN2, or GW5074; CQ was used where indicated to test autophagy dependence.

### *ATXN3* repeat analysis from SCA3 cohort

For *ATXN3* repeat analysis, expanded and normal alleles from 1303 patients with SCA3 were summarized separately by CAG-repeat range and frequency distribution. The proportion of normal alleles carrying 27 or fewer CAG repeats was calculated.

### Statistical analysis

Statistical analyses were performed using GraphPad Prism 9.5 and R. Data are shown as mean ± SD or mean ± SEM, as indicated in the figure legends. Sample sizes, statistical tests, and multiple-comparison procedures are specified in the corresponding legends. Comparisons between two groups were performed using two-tailed Student’s *t* test. Comparisons among three or more groups were performed using one-way ANOVA followed by Dunnett’s multiple-comparisons test unless otherwise indicated. Welch’s one-way ANOVA followed by Dunnett’s T3 multiple-comparisons test was used when variances were unequal. Experiments involving two factors were analyzed by two-way ANOVA followed by Sidak’s or Dunnett’s multiple-comparisons test, as indicated in the figure legends. Time-course behavioral data were analyzed by two-way repeated-measures ANOVA, with treatment as the between-subject factor and time as the within-subject factor. The Geisser-Greenhouse correction was applied to effects involving the repeated-measures factor. Prespecified overall treatment comparisons across all time points were performed versus the DMSO control using Dunnett-adjusted multiple comparisons. Repeated-measures analyses were performed in R, whereas other statistical analyses and graphing were performed using GraphPad Prism 9.5. *P* < 0.05 was considered statistically significant. Exact *P* values are shown where applicable.

## Acknowledgements

The authors thank all of the participants for their involvement in this study.

## Author Contributions

Study design and conceptualization: BXL and HJ; Clinical evaluation and data collection: HYY, ZC, CRW, LLW, LLP, LH, HRP and BST; Genetic analysis: HYY, ZC, XKS, and RQ; Biochemical investigations: HYY, ZC, YQG, YT, JWH and NW; Behavioral and histological experiments: HYY, ZC, CRW, JWH and NW; Data analysis and interpretation: HYY, ZC, XKS, LLP, BXL and HJ; Drafting of the manuscript: HYY and ZC; Critical revision of the manuscript for important intellectual content: BST, XYL, and HJ; Funding acquisition: HJ, ZC, LLW, and LLP.

## Data Availability

The datasets generated and analyzed during the current study are available from the corresponding author on reasonable request.

## Ethics approval and consent to participate

Animal experiments were approved by the Animal Welfare Ethics Committee of the Laboratory Animal Department of Central South University and the Animal Ethics Committee of Xiangya Hospital, Central South University (approval numbers CSU-2023-0001 and 2024020019). Human sample collection and cohort analysis were approved by the Medical Ethics Committee of Xiangya Hospital, Central South University (approval numbers 202310206, 2024010125, and 2025030338). Written informed consent was obtained from all participants, and human samples were handled in accordance with the Declaration of Helsinki.

## Abbreviations

ACSF: artificial cerebrospinal fluid;
ATTECs: autophagosome-tethering compounds; BafA1, bafilomycin A1;
CQ: chloroquine;
DMSO: dimethyl sulfoxide; FBS, fetal bovine serum; ICV, intracerebroventricular;
iPSCs: induced pluripotent stem cells; polyQ, polyglutamine;
SCA3: spinocerebellar ataxia type 3; TG, transgenic;
WT: wild-type;

