## Supplemental Table 1 for "Autophagosome-tethering compounds selectively lower mutant ataxin-3 in preclinical models of spinocerebellar ataxia type 3"

**Supplementary Table 1. Antibodies used in this study**

| Antibody | Application | Company | Catalog no. | Dilution |
|---|---|---|---|---|
| Ataxin-3 /ATXN3 | WB | Sigma-Aldrich | MAB5360 | 1:1000 |
| GAPDH | WB | OriGene | TA802519 | 1:2000 |
| β-tubulin | WB | Proteintech | 10094-1-AP | 1:2000 |
| FLAG tag | WB | Proteintech | 20543-1-AP | 1:20000 |
| polyQ (1C2) | WB | Sigma-Aldrich | MAB1574 | 1:1000 |
| GFP tag | WB | Proteintech | 66002-1-Ig | 1:20000 |
| HA tag | WB | Cell Signaling Technology | 3724 | 1:2000 |
| LC3B | WB | Abcam | ab48394 | 1:1000 |
| p62/SQSTM1 | WB | Sigma-Aldrich | P0067 | 1:2000 |
| Ku86 | WB | Proteintech | 16389-1-AP | 1:2000 |
| Histone H3 | WB | Proteintech | 17168-1-AP | 1:2000 |
| ATG5 | WB | Proteintech | 10181-2-AP | 1:1000 |
| Ataxin-2 /ATXN2 | WB | Proteintech | 21776-1-AP | 1:1000 |
| HTT | WB | Sigma-Aldrich | MAB2166 | 1:1000 |
| THAP11 | WB | Proteintech | 23030-1-AP | 1:1000 |
| β-actin | WB | Proteintech | 66009-1-Ig | 1:20000 |
| Streptavidin-HRP | Streptavidin blot | GenScript | M00091 | 1:5000 |
| Goat anti-Mouse IgG, HRP | WB secondary | Invitrogen | 31430 | 1:10000 |
| Goat anti-Rabbit IgG, HRP | WB secondary | Invitrogen | 31460 | 1:10000 |
| Ataxin-3 /ATXN3 | IHC | Sigma-Aldrich | MAB5360 | 1:2000 |
| Calbindin | IHC | Proteintech | 14479-1-AP | 1:2000 |
| Goat anti-Mouse IgG, HRP | IHC secondary | Servicebio | G1214 | 1:200 |
| Goat anti-Rabbit IgG, HRP | IHC secondary | Servicebio | G1213 | 1:200 |

| | | | | |
|---|---|---|---|---|
| GFAP | IF | Proteintech | 16825-1-AP | 1:200 |
| IBA1 | IF | Abcam | ab283319 | 1:100 |
| OCT4 | IF | Cell Signaling Technology | 2750 | 1:200 |
| SOX2 | IF | Abcam | ab97959 | 1:500 |
| SSEA4 | IF | Cell Signaling Technology | 4755 | 1:200 |
| NANOG | IF | Cell Signaling Technology | 4903 | 1:200 |
| β3-tubulin/TUJ1 | IF | Proteintech | 66240-1-Ig | 1:200 |
| MAP2 | IF | Abcam | ab32454 | 1:200 |
| CoraLite488-conjugated anti-Mouse IgG | IF secondary | Proteintech | SA00013-1 | 1:500 |
| CoraLite488-conjugated anti-Rabbit IgG | IF secondary | Proteintech | SA00013-2 | 1:500 |
| CoraLite594-conjugated anti-Mouse IgG | IF secondary | Proteintech | SA00013-3 | 1:500 |
| CoraLite594-conjugated anti-Rabbit IgG | IF secondary | Proteintech | SA00013-4 | 1:500 |

Notes: WB, western blotting; IHC, immunohistochemistry; IF, immunofluorescence. The anti-ATXN3 antibody MAB5360 is listed separately for WB and IHC because different dilutions were used.
