## Supplemental Figures for "Autophagosome-tethering compounds selectively lower mutant ataxin-3 in preclinical models of spinocerebellar ataxia type 3"

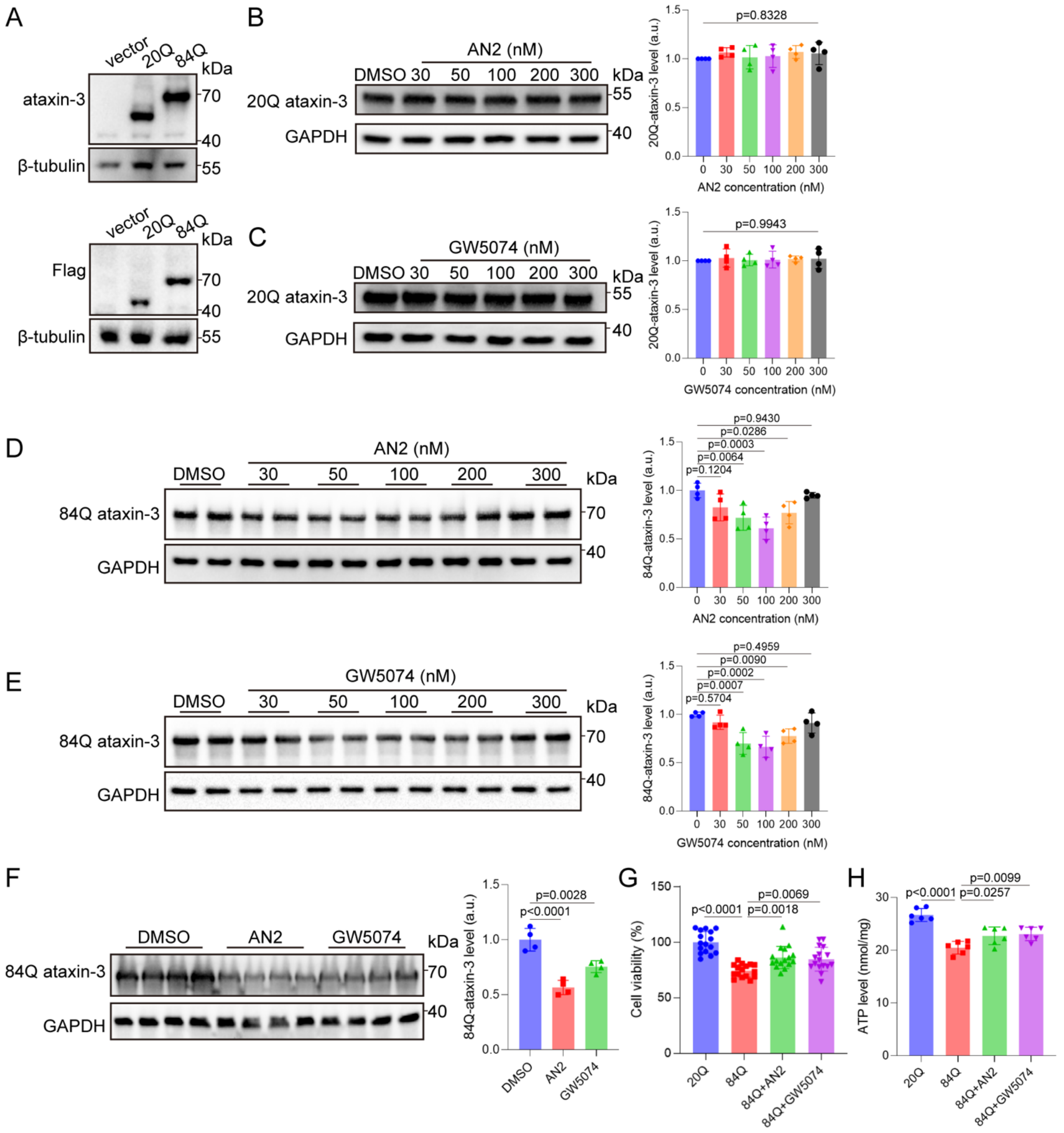


**Supplementary Fig. 1. AN2 and GW5074 selectively reduce mutant ataxin-3 and improve cellular phenotypes in HEK293T models.**

**(A)** Representative immunoblots of HEK293T cells transfected with empty vector, ATXN3-20Q-FLAG, or ATXN3-84Q-FLAG plasmids, showing specific ataxin-3 and FLAG signals. **(B to E)** Representative immunoblots and quantification of 20Q ataxin-3 **(B and C)** or 84Q ataxin-3 **(D and E)** in transiently transfected HEK293T cells treated with DMSO or the indicated concentrations of

AN2 or GW5074 for 48 hours. Protein abundance was normalized to GAPDH and then to the mean value of the DMSO group. Data are shown as mean ± SD; n = 4. **(F)** Representative immunoblot and quantification of 84Q ataxin-3 in transiently transfected HEK293T cells treated with DMSO, AN2 (100nM), or GW5074 (100nM) for 48 hours. Protein abundance was normalized to GAPDH and then to the mean value of the DMSO group. Data are shown as mean ± SD; n = 4. For **(B) to (F)**, statistical analysis was performed by one-way ANOVA followed by Dunnett's multiple-comparisons test versus the DMSO group. **(G)** Cell viability of HEK293T cells expressing 20Q or 84Q ataxin-3 after treatment with DMSO, AN2 (100nM), or GW5074 (100nM). Values were normalized to the mean value of the 20Q ataxin-3 group and expressed as percentages. Data are shown as mean ± SD; n = 16. **(H)** ATP content in HEK293T cells expressing 20Q or 84Q ataxin-3 after treatment with DMSO, AN2 (100nM), or GW5074 (100nM). Data are shown as mean ± SD; n = 6. For **(G)** and **(H)**, statistical analysis was performed by one-way ANOVA followed by Dunnett's multiple-comparisons test versus the 84Q ataxin-3 DMSO group. *P* values are shown in the plots.

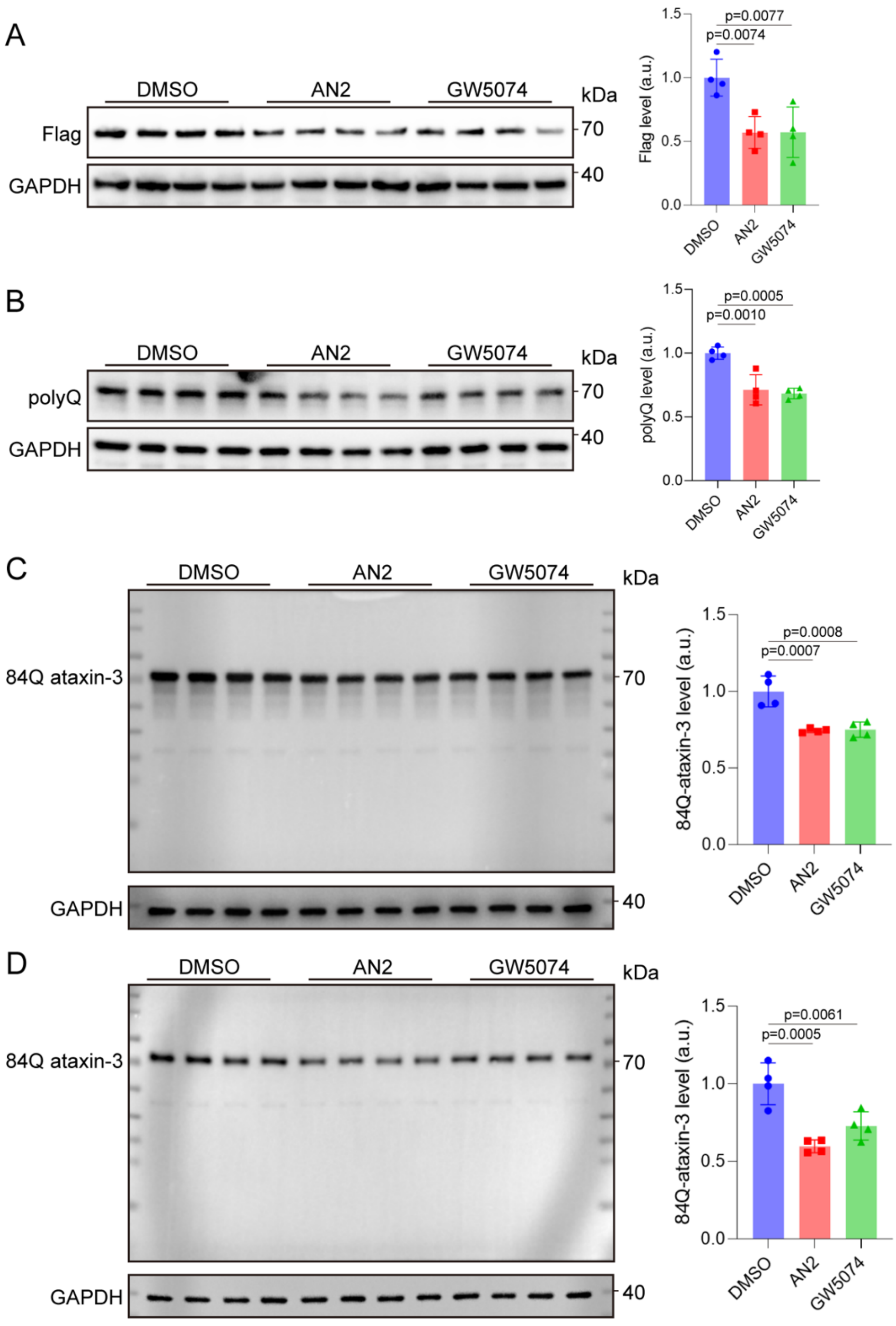


**Supplementary Fig. 2. AN2 and GW5074 reduce 84Q ataxin-3 independently of antibody recognition and without detectable low-molecular-weight ataxin-3 fragments.**

**(A and B)** Representative immunoblots and quantification of 84Q ataxin-3 in HEK293T cells transiently expressing ATXN3-84Q-FLAG after treatment with DMSO, AN2 (100nM), or GW5074 (100nM) for 48 hours. Immunoblots were detected using an anti-FLAG antibody **(A)** or the anti-polyQ antibody 1C2 **(B)**. Protein abundance was normalized to GAPDH and then to the mean value of the DMSO group. **(C and D)** Representative whole-membrane immunoblots and quantification of full-length 84Q ataxin-3 in HEK293T cells transiently expressing ATXN3-84Q-FLAG **(C)** or SH-SY5Y cells stably expressing ATXN3-84Q-FLAG **(D)** after treatment with DMSO, AN2 (100nM), or GW5074 (100nM) for 48 hours. Immunoblots were detected using an anti-ataxin-3 antibody to examine full-length 84Q ataxin-3 and potential low-molecular-weight ataxin-3-reactive fragments. Protein abundance was normalized to GAPDH and then to the mean value of the DMSO group. Data are shown as mean ± SD; n = 4. Statistical analysis was performed by one-way ANOVA followed by Dunnett's multiple-comparisons test versus the DMSO group. *P* values are shown in the plots.

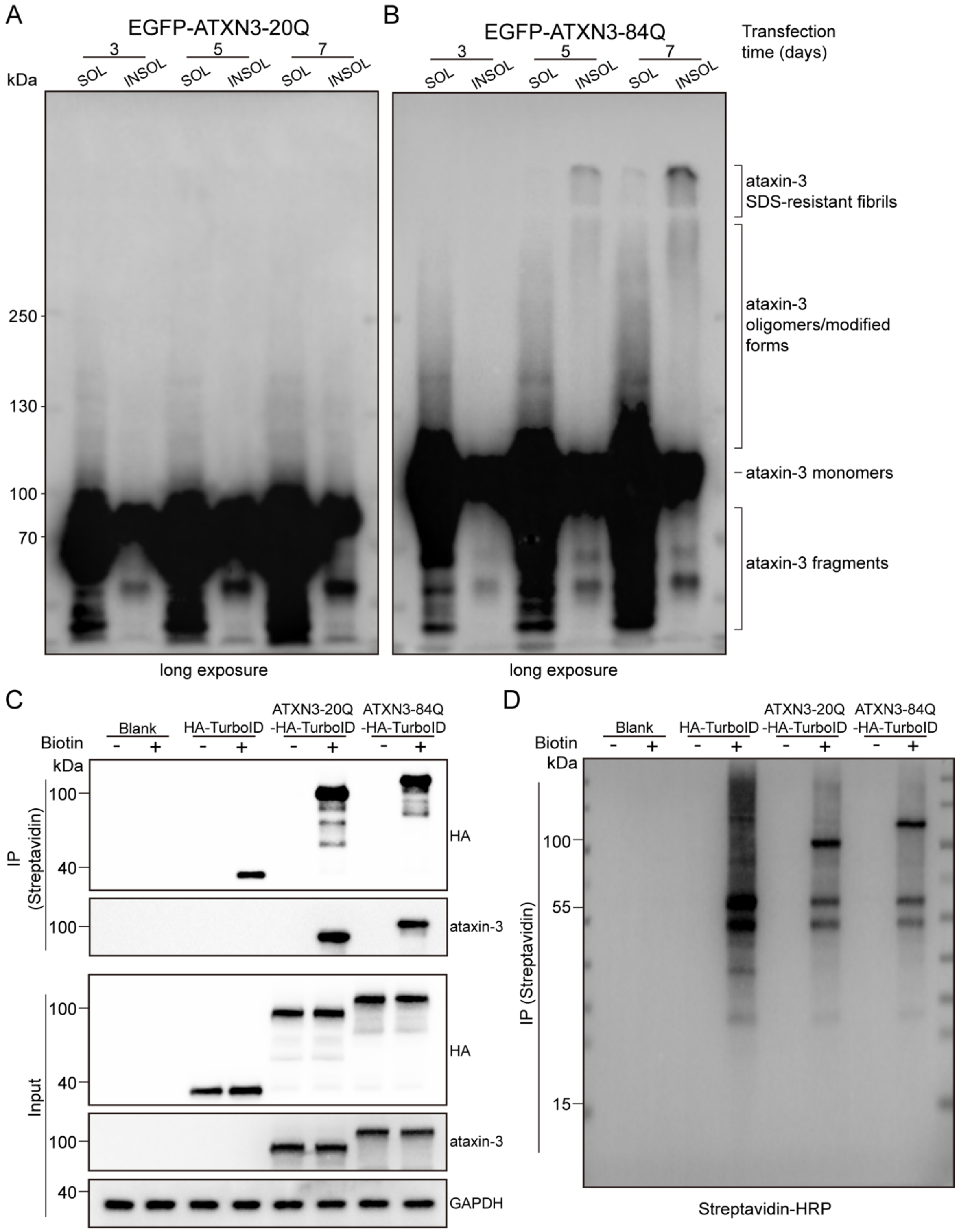


**Supplementary Fig. 3. Validation of SDS-resistant mutant ataxin-3 species and ATXN3-HA-TurboID proximity labeling.**

**(A and B)** Representative long-exposure immunoblots of Triton X-100-soluble and Triton X-100-

insoluble fractions from HEK293T cells expressing EGFP-ATXN3-20Q **(A)** or EGFP-ATXN3-84Q **(B)** at 3, 5, and 7 days after transfection. SDS-resistant ataxin-3 species were detected mainly in the insoluble fractions of EGFP-ATXN3-84Q-expressing cells at days 5 and 7. SOL, Triton X-100-soluble fraction; INSOL, Triton X-100-insoluble fraction. **(C)** Validation of the ATXN3-HA-TurboID proximity-labeling system. HEK293T cells were transfected with HA-TurboID, ATXN3-20Q-HA-TurboID, or ATXN3-84Q-HA-TurboID, and treated with or without 100μM biotin for 4 hours. Whole-cell lysates and streptavidin-enriched fractions were analyzed by immunoblotting for HA and ataxin-3. GAPDH was used as a loading control for input samples. **(D)** Streptavidin-HRP blot of streptavidin-enriched fractions from the same experimental conditions as in **(C)**, showing biotin-dependent proximity labeling by TurboID fusion proteins.

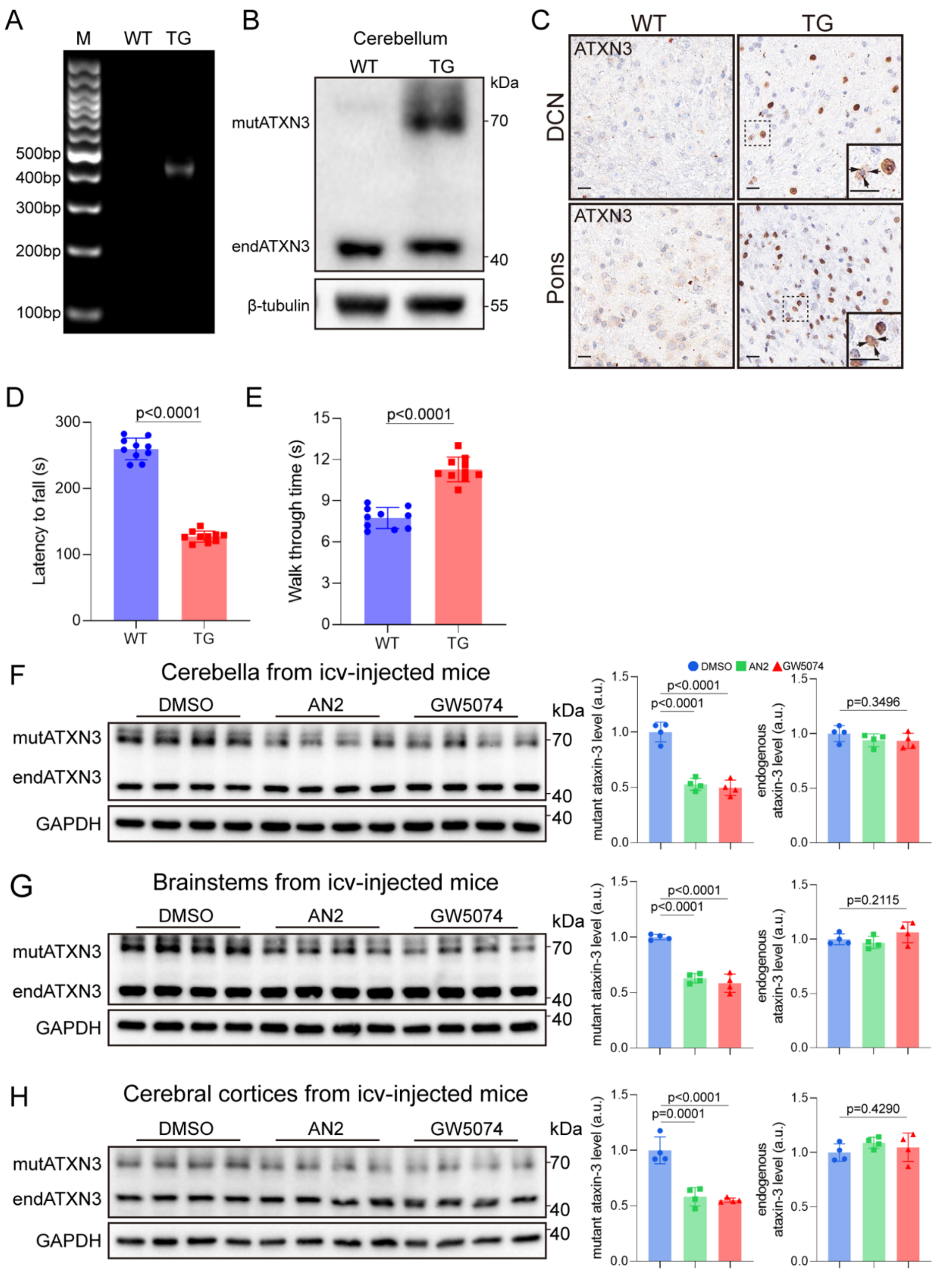


**Supplementary Fig. 4. Characterization of SCA3 transgenic mice and mutant ataxin-3 lowering after intracerebroventricular AN2 or GW5074 administration.**

**(A)** Agarose gel electrophoresis of PCR products for mouse genotyping. A specific 400- to 500-bp amplicon was detected in transgenic (TG) mice but not in wild-type (WT) mice. M, 100-bp DNA ladder. **(B)** Representative immunoblot of ataxin-3 in cerebellar lysates from WT and TG mice.

mutATXN3, human mutant ataxin-3; endATXN3, endogenous mouse ataxin-3. **(C)** Representative immunohistochemical staining for ataxin-3 in the deep cerebellar nuclei (DCN) and pons of WT and TG mice. Insets show higher-magnification views of nuclear ataxin-3-positive inclusions in TG mice. Scale bars, 20μm. **(D to E)** Motor phenotypes of 15-month-old male WT and TG mice. Quantification of latency to fall in the rotarod test **(D)**, time to traverse the balance beam **(E)**. For the rotarod and balance beam tests, each value represents the mean of three trials per mouse. Data are shown as mean ± SD; n = 10. Statistical analysis was performed by two-tailed Student's *t* test. **(F to H)** Representative immunoblots and quantification of ataxin-3 in the cerebellum **(F)**, brainstem **(G)**, and cerebral cortex **(H)** of 8-month-old male TG mice treated by intracerebroventricular injection with DMSO, AN2 (25μM), or GW5074 (25μM) once daily for 10 days. mutATXN3, human mutant ataxin-3; endATXN3, endogenous mouse ataxin-3. Protein abundance was normalized to GAPDH and then to the mean value of the DMSO group. Data are shown as mean ± SD; n = 4. Statistical analysis was performed by one-way ANOVA followed by Dunnett's multiple-comparisons test versus the DMSO group. *P* values are shown in the plots where applicable.

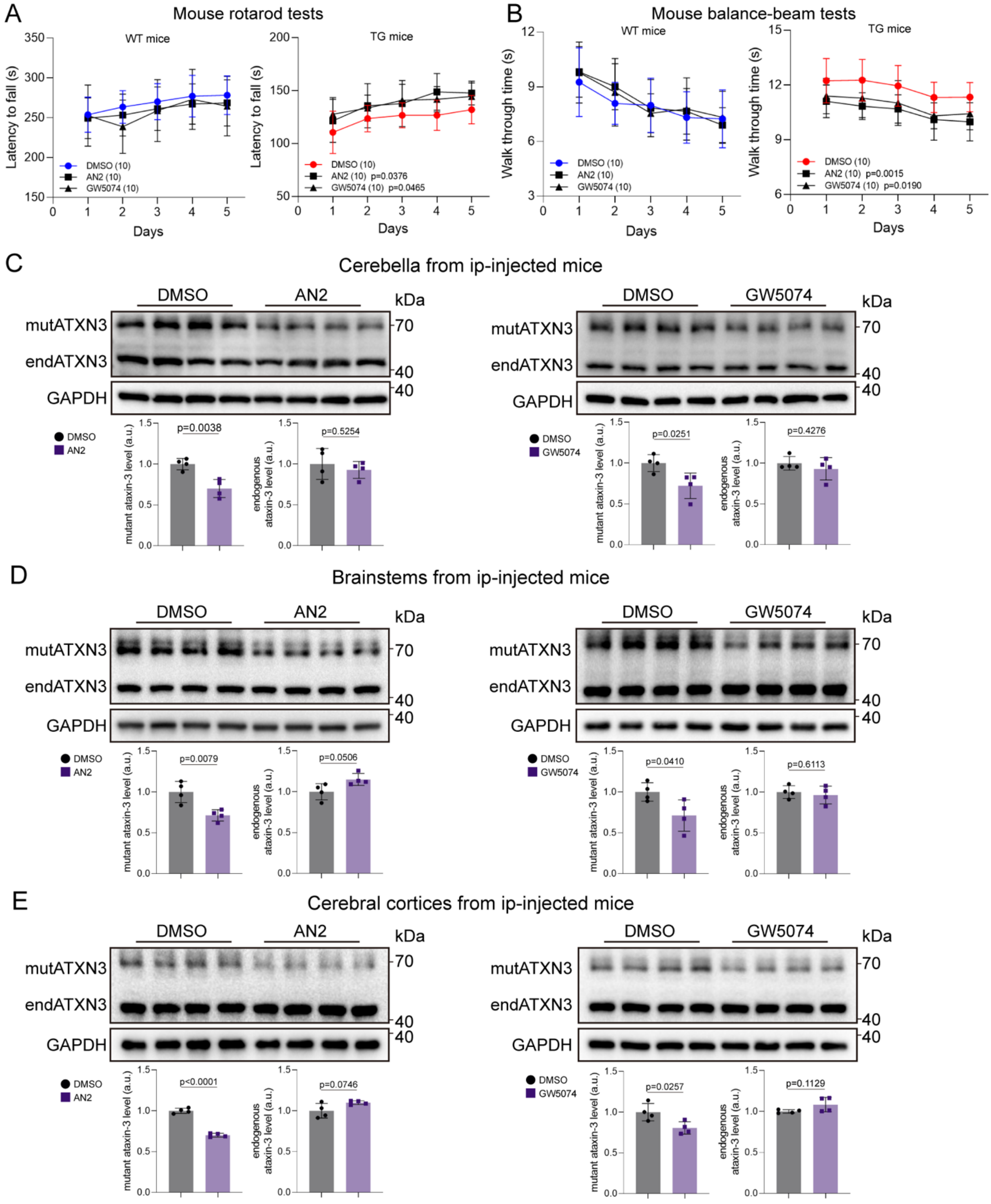


**Supplementary Fig. 5. Intraperitoneal AN2 and GW5074 improve motor phenotypes and reduce mutant ataxin-3 in male SCA3 transgenic mice.**

**(A and B)** Motor performance of 15-month-old male WT and TG mice treated by intraperitoneal injection with DMSO, AN2 (0.5mg/kg), or GW5074 (0.5mg/kg) once daily for 14 days.

Quantification of latency to fall in the rotarod test **(A)** and time to traverse the balance beam **(B)**. For each mouse, the value represents the mean of three trials. Data are shown as mean ± SD; n = 10. Statistical analysis was performed using two-way repeated-measures ANOVA with Dunnett-adjusted comparisons of the overall treatment effects versus DMSO. *P* values shown in the plots represent comparisons across all time points. **(C to E)** Representative immunoblots and quantification of ataxin-3 in the cerebellum **(C)**, brainstem **(D)**, and cerebral cortex **(E)** from 15-month-old male TG mice treated with DMSO, AN2, or GW5074 using the same intraperitoneal dosing schedule. mutATXN3, human mutant ataxin-3; endATXN3, endogenous mouse ataxin-3; i.p., intraperitoneal injection. Protein abundance was normalized to GAPDH and then to the mean value of the corresponding DMSO control. Data are shown as mean ± SD; n = 4. Statistical analysis was performed by two-tailed Student's *t* test comparing each compound group with its corresponding DMSO control. *P* values are shown in the plots.

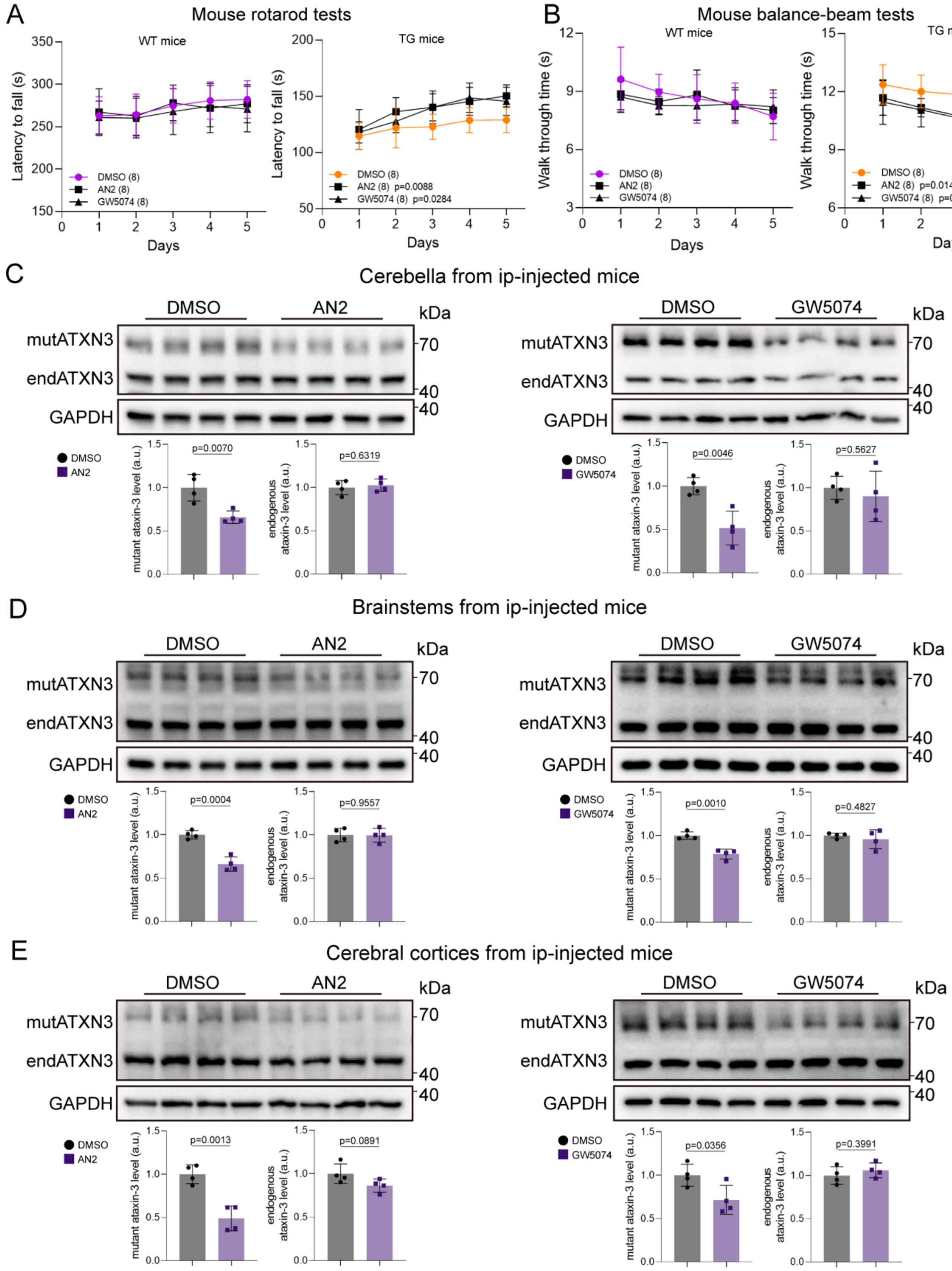


**Supplementary Fig. 6. Intraperitoneal AN2 and GW5074 improve motor phenotypes and reduce mutant ataxin-3 in female SCA3 transgenic mice.**

**(A and B)** Motor performance of 15-month-old female WT and TG mice treated by intraperitoneal injection with DMSO, AN2 (0.5mg/kg), or GW5074 (0.5mg/kg) once daily for 14 days.

Quantification of latency to fall in the rotarod test **(A)** and time to traverse the balance beam **(B)**. For each mouse, the value represents the mean of three trials. Data are shown as mean ± SD; n = 8. Statistical analysis was performed using two-way repeated-measures ANOVA with Dunnett-adjusted comparisons of the overall treatment effects versus DMSO. *P* values shown in the plots represent comparisons across all time points. *P* values shown in the legends indicate the overall treatment effect compared with DMSO. **(C to E)** Representative immunoblots and quantification of ataxin-3 in the cerebellum **(C)**, brainstem **(D)**, and cerebral cortex **(E)** from 15-month-old female TG mice treated with DMSO, AN2, or GW5074 using the same intraperitoneal dosing schedule. mutATXN3, human mutant ataxin-3; endATXN3, endogenous mouse ataxin-3; i.p., intraperitoneal injection. Protein abundance was normalized to GAPDH and then to the mean value of the corresponding DMSO control. Data are shown as mean ± SD; n = 4. Statistical analysis was performed by two-tailed Student's *t* test comparing each compound group with its corresponding DMSO control. *P* values are shown in the plots.

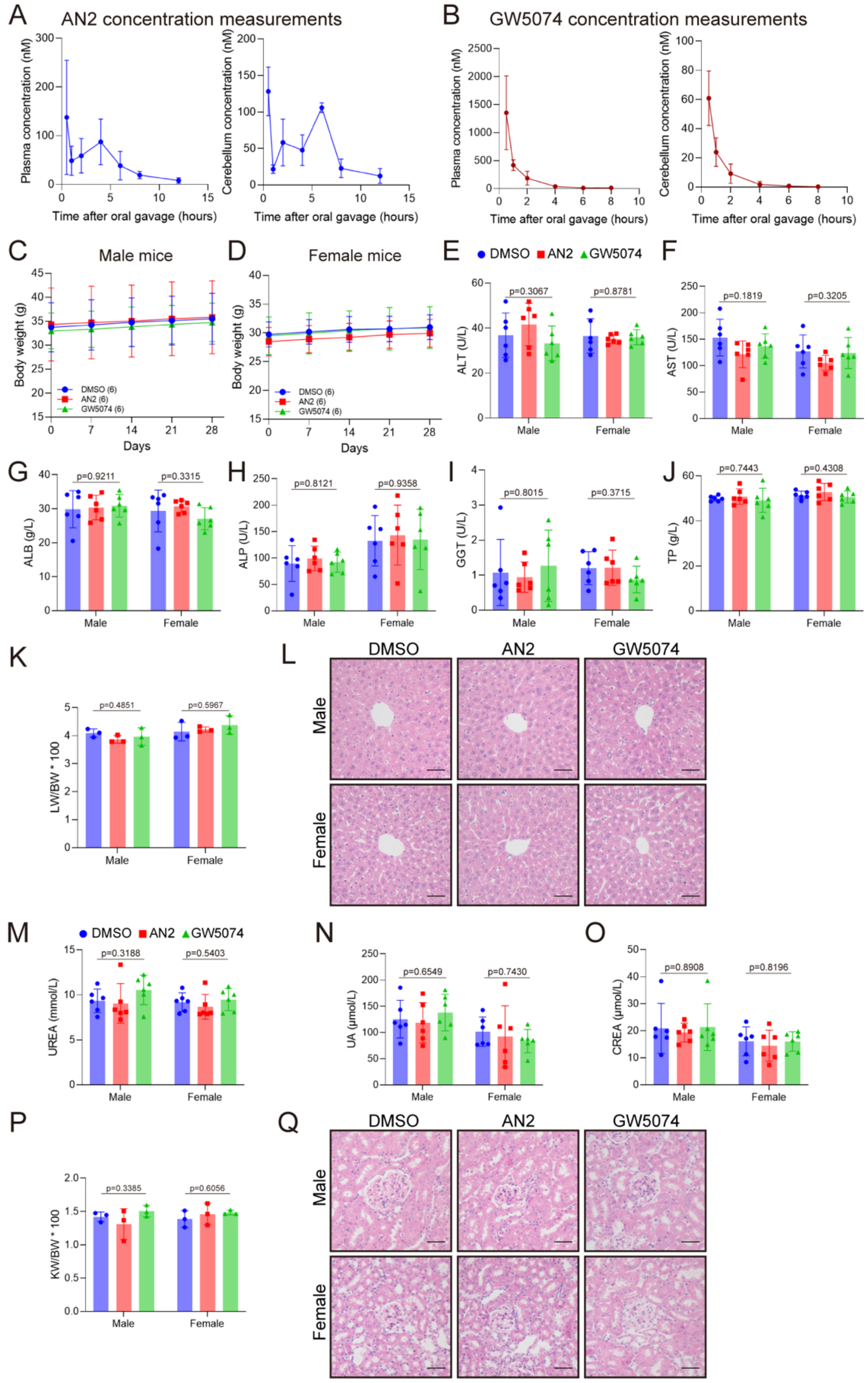


**Supplementary Fig. 7. Oral exposure and safety assessment of AN2 and GW5074 in wild-type mice.**

**(A and B)** Plasma and cerebellar concentrations of AN2 **(A)** and GW5074 **(B)** at the indicated time points after a single oral gavage dose of 15mg/kg, measured by liquid chromatography-mass spectrometry. Data are shown as mean ± SD; n = 3 to 4. **(C and D)** Body weight curves of 15-month-old male **(C)** and female **(D)** wild-type mice orally treated with DMSO, AN2 (15mg/kg), or GW5074 (15mg/kg) for 28 days. Data are shown as mean ± SD; n = 6. Statistical analysis was performed by two-way repeated-measures ANOVA. **(E to J)** Serum liver-function-related biochemical indices after 28 days of oral treatment, including ALT **(E)**, AST **(F)**, ALB **(G)**, ALP **(H)**, GGT **(I)**, and TP **(J)**. Data are shown as mean ± SD; n = 6. **(K)** Liver weight-to-body weight ratio (LW/BW) after 28 days of oral treatment. Data are shown as mean ± SD; n = 3. **(L)** Representative hematoxylin and eosin staining of liver sections from male and female mice after 28 days of oral treatment. Scale bars, 50μm. **(M to O)** Serum kidney-function-related biochemical indices after 28 days of oral treatment, including UREA **(M)**, UA **(N)**, and CREA **(O)**. Data are shown as mean ± SD; n = 6. **(P)** Kidney weight-to-body weight ratio (KW/BW) after 28 days of oral treatment. Data are shown as mean ± SD; n = 3. **(Q)** Representative hematoxylin and eosin staining of kidney sections from male and female mice after 28 days of oral treatment. Scale bars, 50μm. For **(E) to (K)** and **(M) to (P)**, statistical analysis was performed separately within each sex by one-way ANOVA.

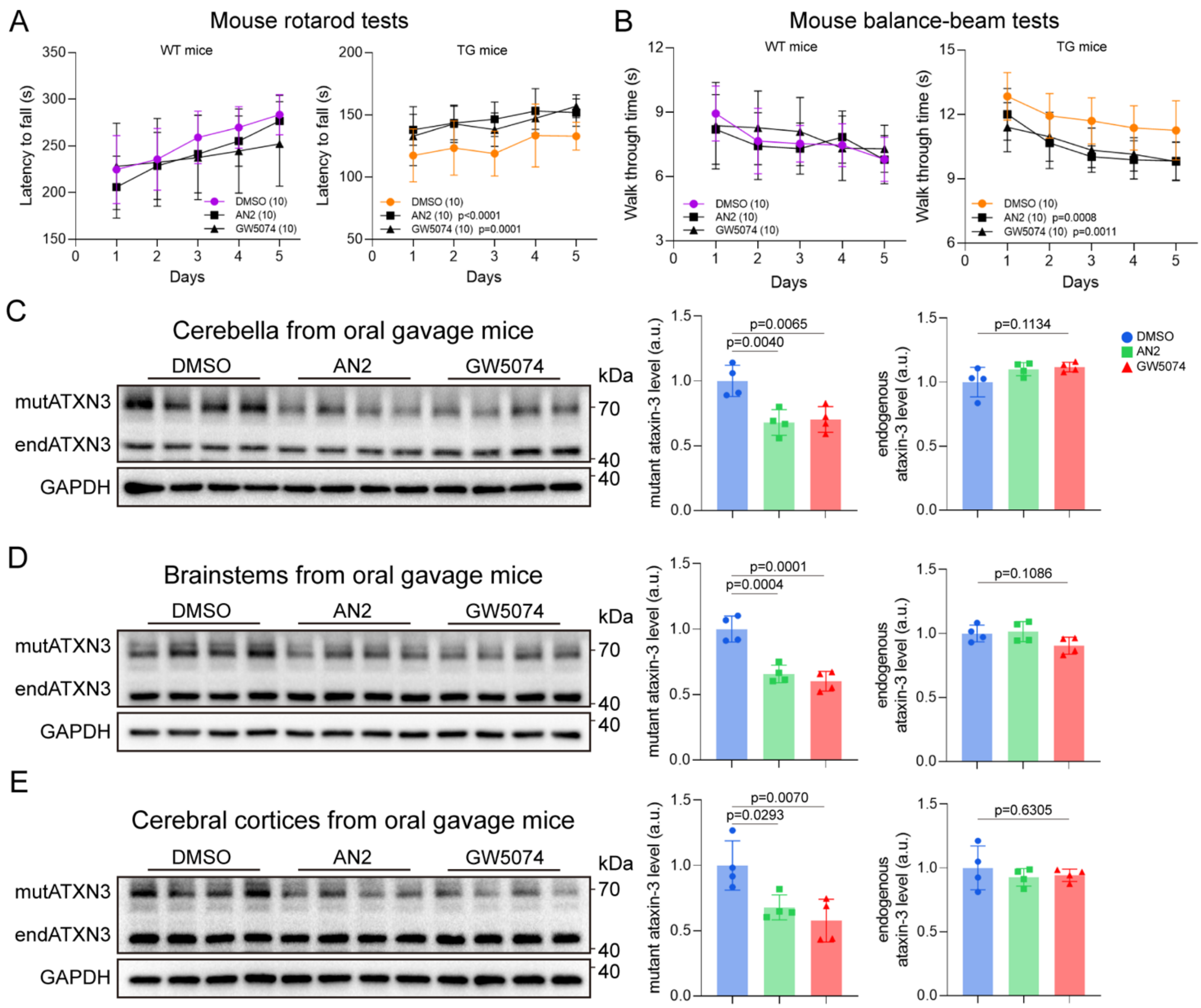


**Supplementary Fig. 8. Oral AN2 and GW5074 improve motor phenotypes and reduce mutant ataxin-3 in female SCA3 transgenic mice.**

Fifteen-month-old female wild-type (WT) or SCA3 transgenic (TG) mice in the indicated experimental groups were orally treated with DMSO, AN2 (15mg/kg), or GW5074 (15mg/kg) for 28 days. **(A and B)** Rotarod performance **(A)** and balance-beam performance **(B)** of WT and TG mice after oral treatment. Rotarod performance is shown as latency to fall, and balance-beam performance is shown as traversal time. For each mouse, the value represents the mean of three trials. Data are shown as mean ± SD; n = 10. Statistical analysis was performed using two-way repeated-measures ANOVA with Dunnett-adjusted comparisons of the overall treatment effects versus DMSO. *P* values shown in the plots represent comparisons across all time points. The *P* values shown in the legends indicate the overall treatment effects compared with DMSO. **(C to E)** Representative immunoblots and quantification of mutant human ataxin-3 and endogenous mouse ataxin-3 in the cerebellum **(C)**, brainstem **(D)**, and cerebral cortex **(E)** of treated female TG mice. Protein abundance was normalized to GAPDH and then to the mean value of the DMSO group. Data are shown as mean ± SD; n = 4. Statistical analysis was performed by one-way ANOVA followed by Dunnett's multiple-comparisons test versus the DMSO group. mutATXN3, mutant human ataxin-3; endATXN3, endogenous mouse ataxin-3.

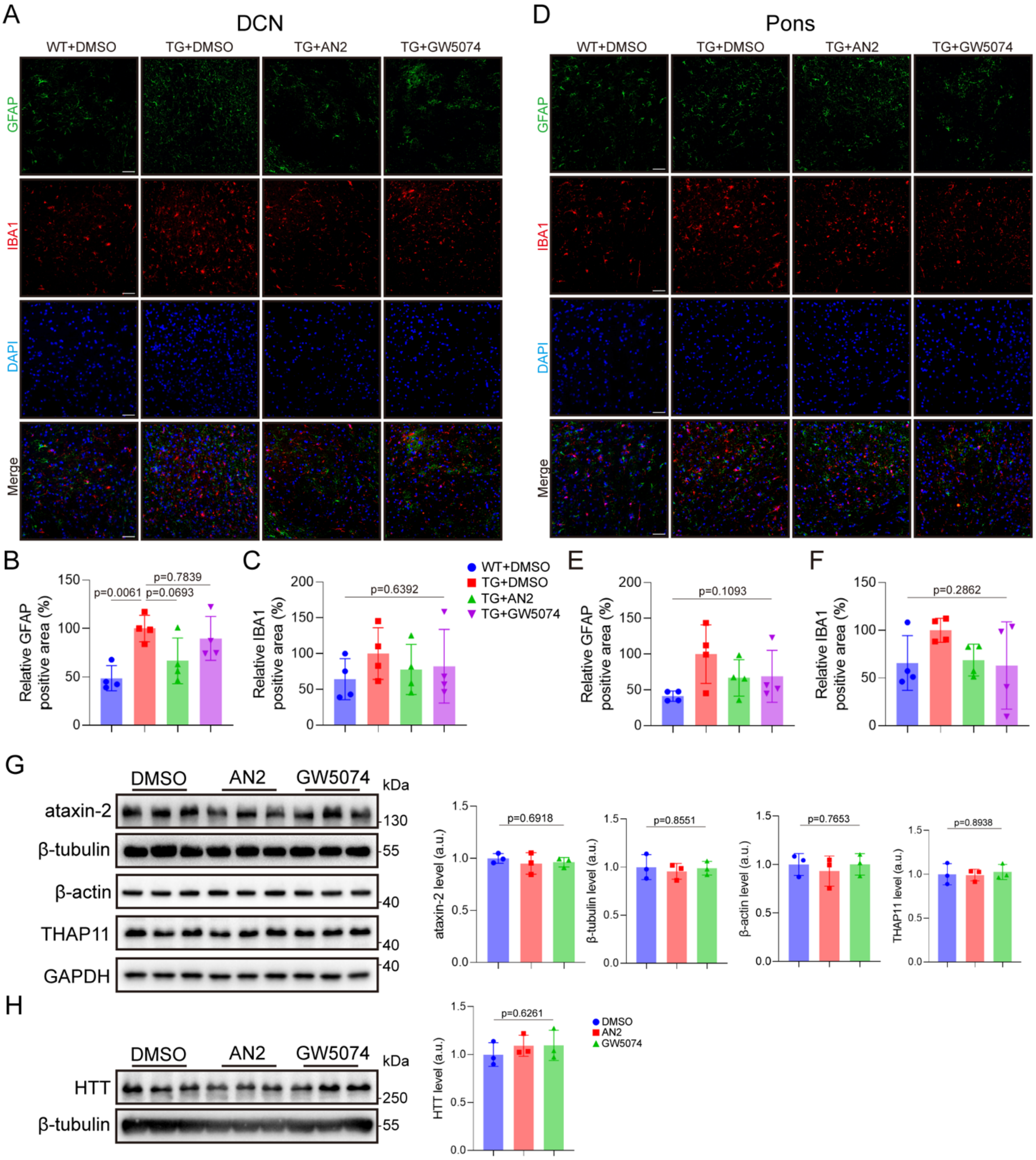


**Supplementary Fig. 9. Effects of oral AN2 and GW5074 on glial markers and selected nontarget proteins in SCA3 transgenic mice.**

Fifteen-month-old WT or SCA3 transgenic TG mice in the indicated experimental groups were orally treated with DMSO, AN2 (15mg/kg), or GW5074 (15mg/kg) for 28 days. **(A)** Representative immunofluorescence images of GFAP and IBA1 in the deep cerebellar nuclei (DCN). Nuclei were counterstained with DAPI. Scale bars, 50μm. **(B and C)** Quantification of the GFAP-positive area

**(B)** and IBA1-positive area **(C)** in the DCN. Values were normalized to the mean value of the TG + DMSO group. **(D)** Representative immunofluorescence images of GFAP and IBA1 in the pons. Nuclei were counterstained with DAPI. Scale bars, 50μm. **(E and F)** Quantification of the GFAP-positive area **(E)** and IBA1-positive area **(F)** in the pons. Values were normalized to the mean value of the TG + DMSO group. For **(B) to (F)**, data are shown as mean ± SD; n = 4. Statistical analysis was performed by one-way ANOVA followed by Dunnett's multiple-comparisons test versus the TG + DMSO group for **(B)** and **(C)**, and by one-way ANOVA for **(E)** and **(F)**. **(G and H)** Representative immunoblots and quantification of selected nontarget proteins in cerebellar lysates from treated TG mice. Ataxin-2, β-tubulin, β-actin, THAP11, and GAPDH are shown in **(G)**, and HTT and β-tubulin are shown in **(H)**. Protein abundance was normalized to GAPDH **(G)** or β-tubulin **(H)** and then to the mean value of the DMSO group. Data are shown as mean ± SD; n = 3. Statistical analysis was performed by one-way ANOVA.

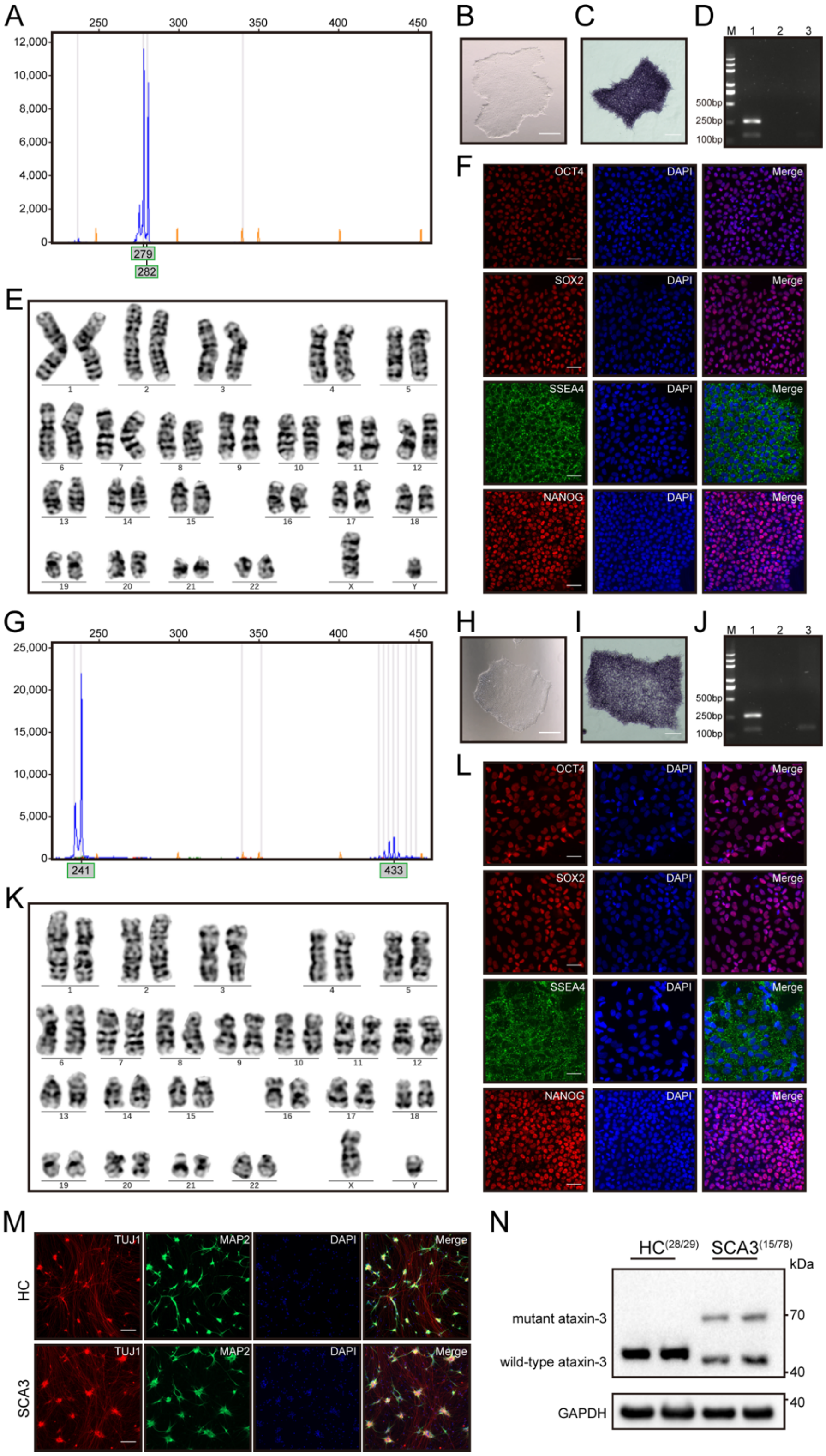


**Supplementary Fig. 10. Characterization of control and SCA3 patient-derived iPSCs and their differentiation into cortical neurons.**

**(A)** Capillary electropherogram showing *ATXN3* CAG-repeat analysis of iPSCs generated from peripheral blood mononuclear cells of the healthy control donor (HC-iPSCs). **(B)** Representative phase-contrast image of HC-iPSC colonies. Scale bar, 100µm. **(C)** Alkaline phosphatase staining of HC-iPSCs. Scale bar, 100µm. **(D)** Agarose gel electrophoresis of PCR-based mycoplasma testing in HC-iPSCs. Lane M, DL2000 DNA marker; lane 1, positive control; lane 2, negative control; lane 3, HC-iPSC sample. **(E)** Karyotype analysis of HC-iPSCs. **(F)** Immunofluorescence staining of pluripotency markers in HC-iPSCs. OCT4, SOX2, and NANOG are shown in red; SSEA4 is shown in green; nuclei were counterstained with DAPI in blue. Scale bars, 50µm. **(G)** Capillary electropherogram showing *ATXN3* CAG-repeat analysis of iPSCs generated from peripheral blood mononuclear cells of the patient with SCA3 (SCA3-iPSCs). **(H)** Representative phase-contrast image of SCA3-iPSC colonies. Scale bar, 100µm. **(I)** Alkaline phosphatase staining of SCA3-iPSCs. Scale bar, 100µm. **(J)** Agarose gel electrophoresis of PCR-based mycoplasma testing in SCA3-iPSCs. Lane M, DL2000 DNA marker; lane 1, positive control; lane 2, negative control; lane 3, SCA3-iPSC sample. **(K)** Karyotype analysis of SCA3-iPSCs. **(L)** Immunofluorescence staining of pluripotency markers in SCA3-iPSCs. OCT4, SOX2, and NANOG are shown in red; SSEA4 is shown in green; nuclei were counterstained with DAPI in blue. Scale bars, 50µm. **(M)** Representative immunofluorescence images of cortical neuron marker expression at day 14 of differentiation from HC-iPSCs and SCA3-iPSCs. β3-tubulin/TUJ1 is shown in red, MAP2 is shown in green, and nuclei were counterstained with DAPI in blue. Scale bars, 100µm. **(N)** Representative immunoblot showing wild-type and mutant ataxin-3 expression in differentiated neurons derived from HC-iPSCs and SCA3-iPSCs. GAPDH was used as a loading control.

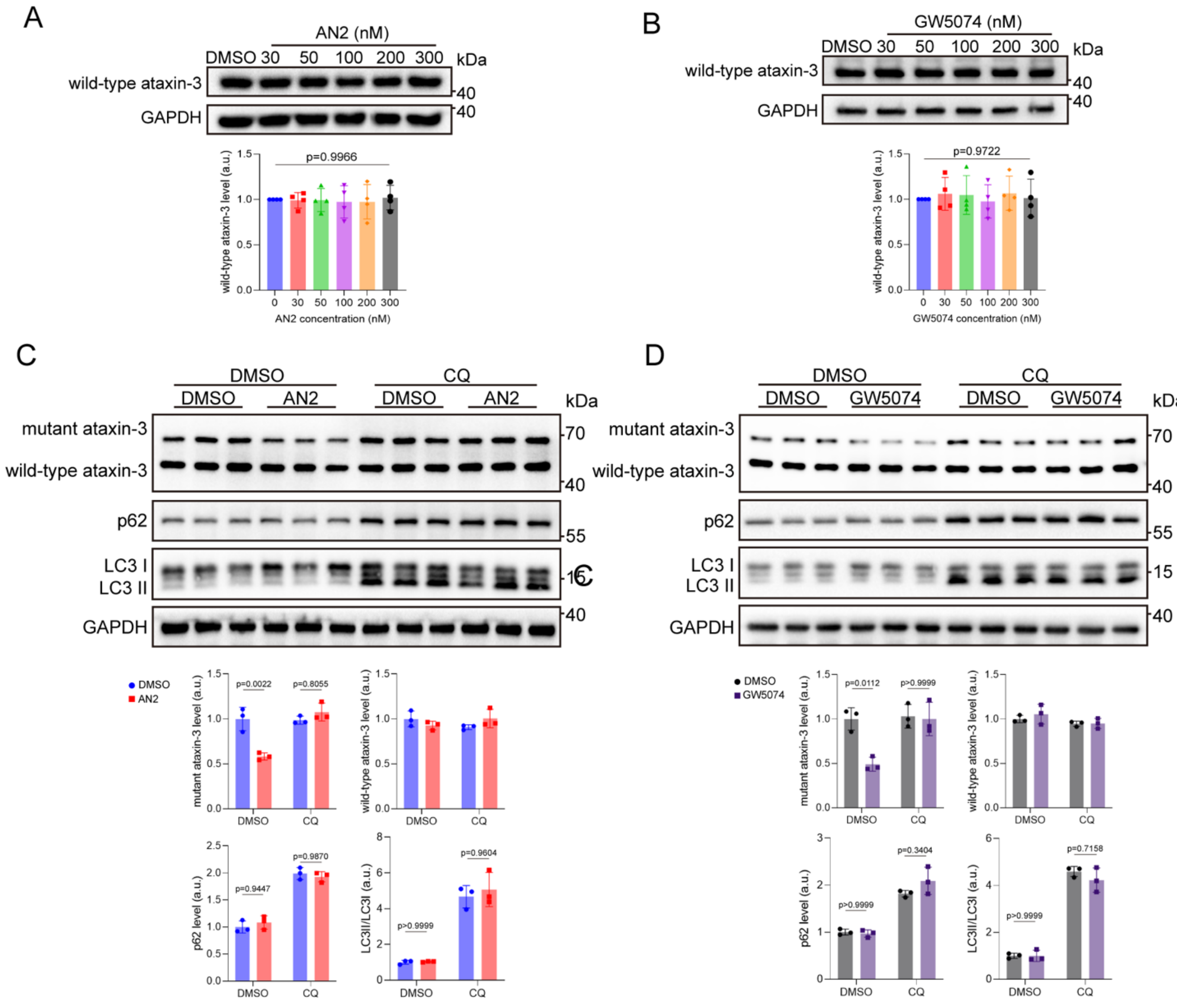


**Supplementary Fig. 11. AN2 and GW5074 spare normal ataxin-3 and require autophagy for mutant ataxin-3 lowering in iPSC-derived neurons.**

**(A and B)** Representative immunoblots and quantification of wild-type ataxin-3 in cortical neurons differentiated from HC-iPSCs after treatment with DMSO or the indicated concentrations of AN2 **(A)** or GW5074 **(B)**. Protein abundance was normalized to GAPDH and then to the mean value of the DMSO group. Data are shown as mean ± SD; n = 4. Statistical analysis was performed by one-way ANOVA. **(C and D)** Representative immunoblots and quantification of mutant ataxin-3, wild-type ataxin-3, p62, and the LC3-II/LC3-I ratio in cortical neurons differentiated from SCA3-iPSCs after pretreatment with or without CQ (40μM) for 12 hours, followed by treatment with DMSO, AN2 (100nM) **(C)**, or GW5074 (100nM) **(D)** until differentiation day 5. Protein abundance was normalized to GAPDH, and values were then normalized to the mean value of the DMSO group without CQ pretreatment. Data are shown as mean ± SD; n = 3. Statistical analysis was performed by two-way ANOVA followed by Sidak's multiple-comparisons test comparing DMSO versus compound treatment in the absence of CQ and the corresponding DMSO versus compound treatment in the presence of CQ. For mutant ataxin-3, the interaction *P* value between CQ pretreatment and compound treatment was 0.0010 for AN2 and 0.0164 for GW5074. For p62 and the LC3-II/LC3-I ratio, the *P* values shown in the plots indicate the corresponding DMSO-versus-compound comparisons under each CQ condition.
